# Interrogation of Enhancer Function by Enhanced CRISPR Epigenetic Editing

**DOI:** 10.1101/761247

**Authors:** Kailong Li, Yuxuan Liu, Hui Cao, Yuannyu Zhang, Zhimin Gu, Xin Liu, Andy Yu, Pranita Kaphle, Kathryn E. Dickerson, Min Ni, Jian Xu

## Abstract

Tissue-specific gene expression requires coordinated control of gene-proximal and -distal *cis*-regulatory elements (CREs), yet functional analysis of gene-distal CREs such as enhancers remains challenging. Here we describe enhanced CRISPR/dCas9-based epigenetic editing systems, enCRISPRa and enCRISPRi, for multiplexed analysis of enhancer function *in situ* and *in vivo*. Using dual effectors capable of re-writing enhancer-associated chromatin modifications, we show that enCRISPRa and enCRISPRi modulate gene transcription by remodeling local epigenetic landscapes at sgRNA-targeted enhancers and associated genes. Comparing with existing methods, the new systems display more robust perturbation of enhancer activity and gene transcription with minimal off-targets. Allele-specific targeting of enCRISPRa to oncogenic *TAL1* super-enhancer modulates *TAL1* expression and cancer progression in xenotransplants. Multiplexed perturbations of lineage-specific enhancers using an enCRISPRi knock-in mouse establish *in vivo* evidence for lineage-restricted essentiality of developmental enhancers during hematopoietic lineage specification. Hence, enhanced CRSIPR epigenetic editing provides opportunities for interrogating enhancer function in native biological contexts.

## INTRODUCTION

Mammalian gene expression requires precisely regulated gene-proximal promoters and gene-distal *cis*-regulatory elements (CREs) such as transcriptional enhancers. Systematic annotation of human epigenomes has identified millions of putative CREs using correlative features such as chromatin accessibility and histone modifications ^1-4^; however, the *in vivo* functions of the vast majority of these elements within their native chromatin remain unknown. This is in part because existing technologies often measure enhancer activity in heterologous assays without native chromatin, and because findings from these assays have not been causally connected with specific target genes or cellular functions during development.

Enhancers are *cis*-regulatory DNA sequences that are bound and regulated by transcription factors (TFs) and chromatin regulators in a highly tissue-specific manner. Putative enhancers are operationally identified using epigenetic signatures including chromatin accessibility (DNase I hypersensitivity or ATAC-seq) and histone marks (H3K4me1/2 and H3K27ac) ^5-7^. Unlike gene-proximal promoters, enhancers can regulate gene transcription over long distances in an orientation-independent and cell-type-specific manner ^8^. As such, fundamental challenges have limited the application of existing technologies in functional analysis of a specific enhancer in a mammalian genome. Reporter assays have historically been used to examine enhancer activity in heterologous cell models ^8^. When combined with high-throughput sequencing, massively parallel reporter assays allow for quantitative analysis of the transcriptional activity of thousands of enhancers in particular cell types ^9, 10^. Together with transgenics, *in vivo* enhancer reporter assays enable evaluation of enhancer function during mammalian development ^11^. These are powerful approaches for assaying TF-mediated transcriptional activity at enhancer DNA sequences, but they have some important limitations including the lack of local chromatin contexts and epigenetic features in heterologous assays, the use of a general promoter such as SV40 rather than the enhancer’s endogenous promoter, the inability to identify the target genes of enhancers, and the inadequacy to model combinatorial regulation by multiple enhancers at native chromatin. Additionally, conventional gene targeting or genome editing approaches have been utilized to knockout (KO) or mutate specific enhancers in cell lines or animal models ^12, 13^; however, they require genetic engineering which remains low-throughput and laborious. Furthermore, high-resolution saturating screens of *cis*-regulatory elements rely on loss-of-function and do not permit gain-of-function analyses ^14-16^.

Recently, major advances have been made in the modulation of endogenous gene expression by repurposing the CRISPR/Cas9 system ^17-29^. By coupling the deactivated Cas9 (dCas9) to various activator (e.g. VP64 ^18-20^, p300 ^22, 28^, and SAM ^21^) or repressor (e.g. KRAB ^18, 19, 23, 28^, LSD1 ^29^, and DNMT3A/3L ^26^) domains, transcriptional perturbation of specific genes were achieved. While the most commonly used dCas9 activator or repressor complexes such as dCas9-VP64, dCas9-SAM or dCas9-KRAB can effectively modulate transcription when tethered to gene-proximal promoters, the effect declines rapidly when its target region moves away from proximal promoter sequences ^18-21^. This is likely because VP64 or KRAB preferentially interferes with the basal transcription initiation and/or elongation apparatus operating at gene promoters ^29, 30^. Since distal CREs such as enhancers may not rely on the basal transcription apparatus, these methods were ineffective and variable in modulating enhancer activity ^18-21^.

Here we develop enhanced CRISPR epigenetic editing systems, enCRISPRa and enCRISPRi, to interrogate enhancer function using dCas9 with dual effectors that specifically modulate epigenetic modifications at enhancers. Using the human β-globin locus control region (LCR), oncogenic *TAL1* super-enhancer (SE), and hematopoietic lineage-specific enhancers as examples, we show that enCRISPRa and enCRISPRi effectively modulate enhancer function *in vitro*, in xenografts and *in vivo*. Enhanced CRISPR epigenetic editing leads to locus-specific epigenetic reprogramming and interference with TF binding. Single or multiplexed *in vivo* enhancer perturbations using an enCRISPRi mouse model reveal lineage-specific requirements of developmental enhancers during hematopoiesis. Hence, the enhanced CRISPR epigenetic editing systems provide opportunities for functional interrogation of enhancers and other CREs in development and disease.

## RESULTS

### Development of an enCRISPRa System for Enhancer Activation

To assess the functional role of gene-distal enhancers, we devised the enhanced dCas9-based epigenetic perturbation systems for targeted modulation of enhancer activity *in situ* and *in vivo*. Specifically, we employed the structure-guided sgRNA design by adding two MS2 hairpins ^21^, which is recognized by the MCP RNA-binding proteins ^31^. For enhancer activation (enCRISPRa; Fig. 1a), we fused dCas9 with the core domain of histone acetyltransferase p300, which catalyzes H3-Lys27 acetylation (H3K27ac) ^32^, together with the MS2-sgRNA sequence to recruit the MCP-VP64 activator domains. Since H3K27ac is the hallmark of active enhancers ^33, 34^, by doxycycline (Dox)-inducible expression of dCas9-p300, sgRNA-MS2 and MCP-VP64, we engineered an enhancer-targeting dual-activator system (Fig. 1a). As a proof-of-principle test of enCRISPRa in modulating CRE activity, we targeted enCRISPRa to several known enhancers or promoters including the *MYOD* enhancer, the *IL1RN* and *OCT4* promoters ^22^ in HEK293T cells (Fig. 1b). Comparing with existing next-generation dCas9-based activation methods such as dxCas9-VPR ^24^, SunTag ^19^ and SAM ^21^, enCRISPRa showed comparable potency on gene activation when targeted to the *IL1RN* and *OCT4* promoters (Fig. 1b). More importantly, enCRISPRa displayed significantly more robust activation of gene transcription compared to other dCas9 activators when targeted to the *MYOD* enhancer (Fig. 1b).

**Figure 1.**
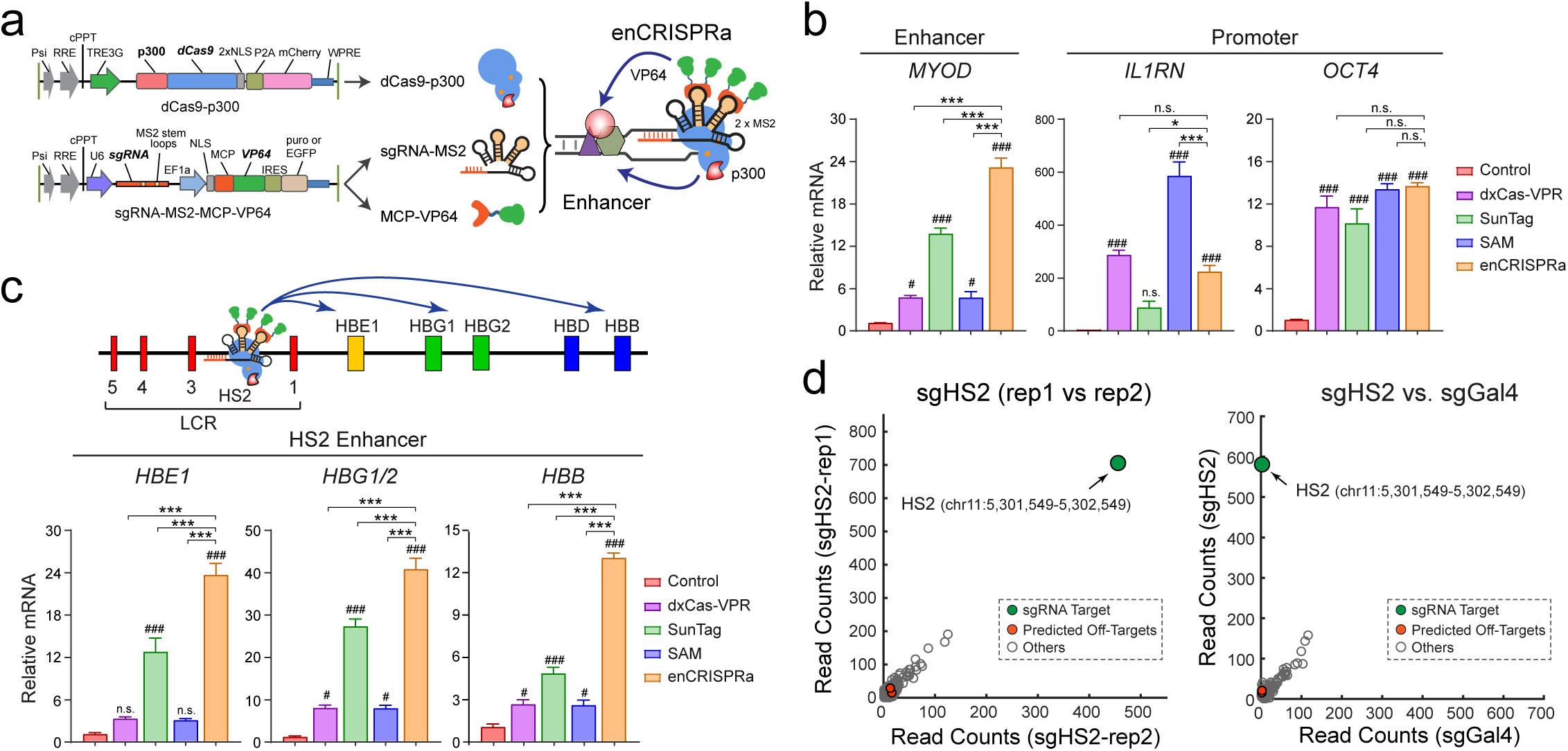
Development of the Dual-Activator enCRISPRa system. **(a)** Schematic of the enCRISPRa system containing three components: a dCas9-p300 fusion protein, the target-specific sgRNA with two MS2 hairpins, and the MCP-VP64 fusion protein. **(b)** Expression of *MYOD, IL1RN* and *OCT4* upon dxCas9-VPR, SunTag, SAM or enCRISPRa-mediated enhancer or promoter activation in HEK293T cells. mRNA expression relative to non-transduced cells is shown as mean ± SEM. The differences between control and dCas9 activators were analyzed by a two-way ANOVA. ^#^*P* < 0.05, ^##^*P* < 0.01, ^###^*P* < 0.001. The difference between enCRISPRa and other dCas9 activators were analyzed by a two-way ANOVA. \**P* < 0.05, \*\**P* < 0.01, \*\*\**P* < 0.001, n.s. not significant. **(c)** Expression of β-globin genes *HBE1, HBG1/2* and *HBB* upon dxCas9-VPR, SunTag, SAM or enCRISPRa-mediated activation of the HS2 enhancer in HEK293T cells. mRNA expression relative to non-transduced cells is shown as mean ± SEM and analyzed by a two-way ANOVA. **(d)** Genome-wide analysis of dCas9 binding in HEK293T cells expressing HS2-specific sgRNA (two replicate ChIP-seq experiments sgHS2-rep1 and sgHS2-rep2) or non-targeting sgGal4. Data points for the sgRNA target regions and the predicted off-targets are shown as *green* and *red*, respectively. The x- and y-axis denote the normalized read counts (*left*) or mean normalized read counts from *N* = 2 independent ChIP-seq experiments (*right*).

We next focused on the well-established HS2 enhancer at the human β-globin locus ^35^ (Fig. 1c). The β-globin locus contains five β-like globin genes (*HBE1, HBG1, HBG2, HBD* and *HBB*) that are developmentally regulated by a shared upstream enhancer cluster or locus control region (LCR) ^36^. The β-globin LCR consists of five discrete enhancers (HS1 to HS5) in which HS2 functions as an erythroid-specific enhancer in transgenic assays ^36^, and provides a paradigm for studying tissue-specific and developmentally regulated gene transcription. We engineered HEK293T cells with Dox-inducible expression of dCas9-p300 or other dCas9 activators ^19, 21, 24^. Upon co-expression of a single HS2-targeting sgRNA, we measured mRNA expression of β-globin genes and compared with existing dCas9 activators. Notably, enCRISPRa-mediated HS2 enhancer activation led to 13.0 to 40.6-fold increases in expression of β-globin genes *HBE1, HBG1/2* and *HBB* relative to the non-transduced controls, which are significantly higher than other dCas9 activation methods (Fig. 1c). Although different dCas9 activators were noted to display variable potencies in gene activation in previous studies ^21, 37-39^, the observed differences could be affected by different cellular contexts, particular target genes, position of sgRNAs (e.g. promoter vs enhancer), transfection conditions, and time to analyze gene expression. Thus, it is important to note that our analyses were performed in the same cell lines using the same sgRNA and transfection protocol, which enabled us to compare the efficacy of different dCas9 activators on enhancer perturbation side-by-side. Moreover, analysis of dCas9 chromatin occupancy in cells co-expressing enCRISPRa and HS2-specific sgRNA (sgHS2) revealed highly reproducible binding at the targeted HS2 enhancer by independent ChIP-seq experiments (Fig. 1d; Table S1). By comparing dCas9 binding in cells expressing sgHS2 or non-targeting sgRNA (sgGal4), we observed highly specific enrichment of dCas9 at HS2 with no additional significant binding at the genome scale (Fig. 1d).

**Development of an enCRISPRi System for Enhancer Repression**

We next devised an enhanced epigenetic editing system for targeted enhancer inhibition (enCRISPRi-LK; Fig. 2a). Specifically, we fused dCas9 with the lysine-specific demethylase LSD1 (or KDM1A), which catalyzes the removal of enhancer-associated H3-Lys4 mono- and dimethylation (H3K4me1/2) ^40^, together with MS2-sgRNA to recruit the MCP-KRAB repressor domains (Fig. 2a). As a proof-of-principle test, we targeted enCRISPRi to the HS2 enhancer in K562 erythroleukemia cells that highly express β-globin genes ^35^. We achieved 3.4 to 13.7-fold repression of β-globin genes (*HBE1, HBG1/2* and *HBB*) by 4 independent sgRNAs (sgHS2-1 to sgHS2-4) relative to non-targeting sgGal4 (Fig. 2b). We further engineered the second version of enCRISPRi using dCas9-KRAB + MCP-LSD1 (enCRISPRi-KL) combination (Fig. 2a). Targeting of either enCRISPRi complex to the HS2 enhancer by 4 independent sgRNAs achieved comparable and significant repression of β-globin genes (Fig. 2b). Comparing with the single effector dCas9-KRAB (K) or dCas9-LSD1 (L) complex, the dual repressor-containing enCRISPRi displayed markedly stronger gene repression (Fig. 2b,c). In addition, enCRISPRi resulted in minimal changes (except β-globin genes) in global transcriptomics by RNA-seq (Fig. 2c). Moreover, ChIP-seq analyses revealed significant enrichment of dCas9 binding at the targeted HS2 enhancer with minimal off-targets by independent replicate experiments and/or comparing to the non-targeting sgGal4 control (Fig. 2d), indicating the locus-specific modulation of target gene transcription.

**Figure 2.**
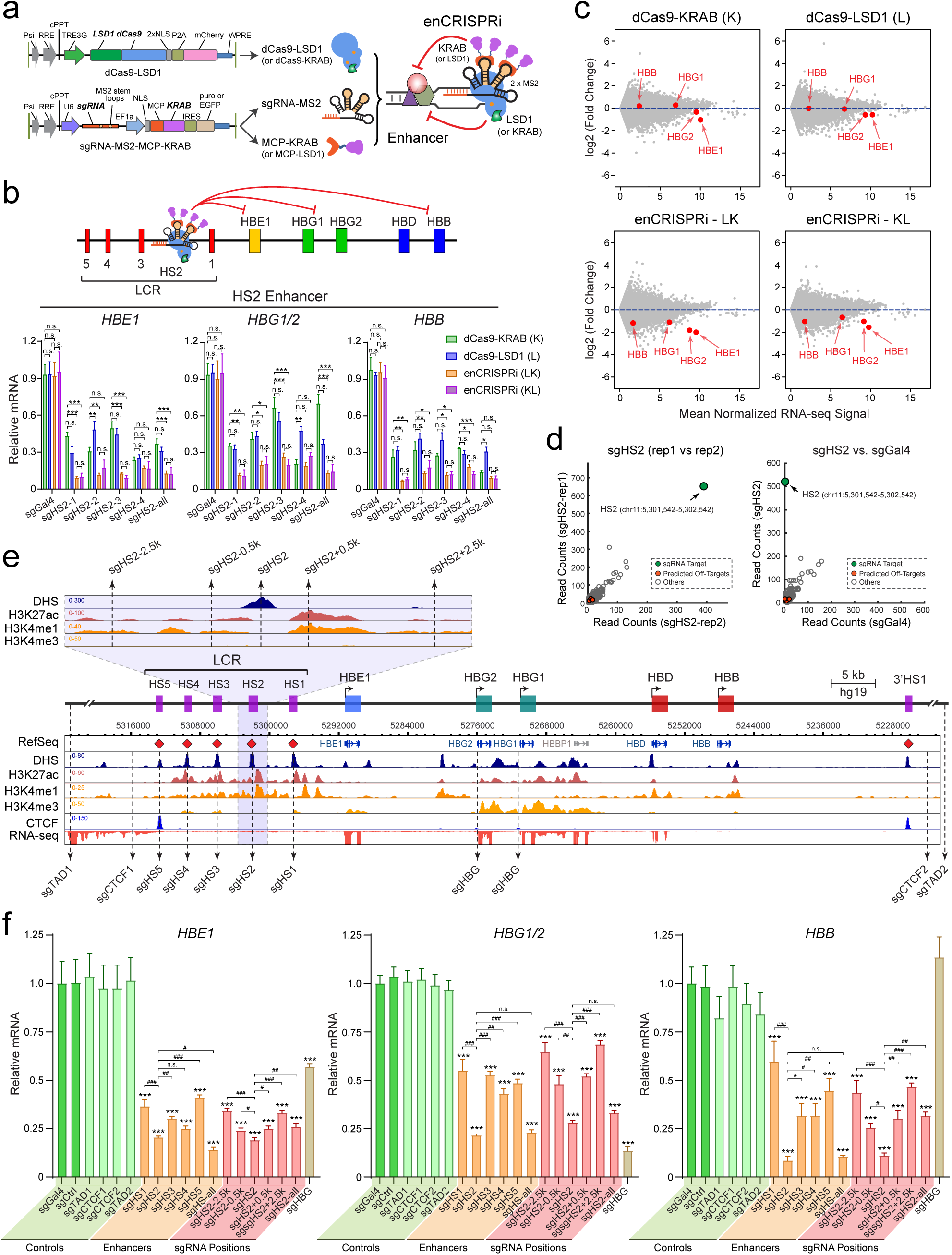
Development of the Dual-Repressor enCRISPRi system. **(a)** Schematic of the enCRISPRi system containing a dCas9-LSD1 fusion protein, the target-specific sgRNA with two MS2 hairpins, and the MCP-KRAB fusion protein. **(b)** Expression of β-globin genes in K562 cells upon dCas9-KRAB (K), dCas9-LSD1 (L) or enCRISPRi (LK and KL)-mediated repression of the HS2 enhancer using four independent HS2-targeting sgRNAs individually (sgHS2-1 to sgHS2-4) or combined (sgHS2-all). mRNA expression relative to non-transduced K562 cells is shown as mean ± SEM and analyzed by a two-way ANOVA. \**P* < 0.05, \*\**P* < 0.01, \*\*\**P* < 0.001, n.s. not significant. **(c)** RNA-seq profiles in K562 cells upon dCas9-KRAB, dCas9-LSD1 or enCRISPRi-mediated repression of the HS2 enhancer using all four HS2-targeting sgRNAs (sgHS2-all). Scatter plot is shown for each gene by the mean of log2 normalized RNA-seq signals as transcripts per million or TPM (*N* = 2 independent experiments) (x-axis) and log2 fold changes of mean TPM in cells expressing sgHS2 and control sgGal4 (y-axis). The β-globin genes (*HBE1, HBG1, HBG2* and *HBB*) are indicated by red arrowheads. **(d)** Genome-wide differential analysis of dCas9 binding in K562 cells expressing HS2-specific sgRNA (sgHS2-rep1 and sgHS2-rep2) or non-targeting sgGal4. Data points for the sgRNA target regions and the predicted off-targets are shown as *green* and *red*, respectively. **(e)** Density maps are shown for DHS, ChIP-seq of H3K27ac, H3K4me1, H3K4me2, CTCF, and RNA-seq at the β-globin cluster (chr11: 5,222,500-5,323,700; hg19). The zoom-in view of the HS2 proximity region is shown on the top. Dashed lines denote the positions of sgRNAs. **(f)** Expression of β-globin genes in K562 cells co-expressing enCRISPRi and target-specific sgRNAs at various positions within the β-globin cluster, control sgRNAs (sgCtrl, sgTAD1, sgTAD2, sgCTCF1 and sgCTCF2) or non-targeting sgGal4. mRNA expression relative to sgGal4 is shown as mean ± SEM. The difference between control sgGal4 and other sgRNAs were analyzed by a two-way ANOVA. \**P* < 0.05, \*\**P* < 0.01, \*\*\**P* < 0.001, n.s. not significant. The differences between sgHS2 and other sgRNAs were analyzed by a two-way ANOVA. ^#^*P* < 0.05, ^##^*P* < 0.01, ^###^*P* < 0.001.

We further explored the effectiveness of enCRISPRi by targeting to single or multiple enhancers at the β-globin LCR enhancer cluster (Fig. 2e). We focused on the enCRISPRi (LK) version given similar effects on gene repression when targeting enCRISPRi LK and KL to the HS2 enhancer (Fig. 2a-c). To this end, we designed sgRNAs for individual LCR enhancers (sgHS1 to sgHS5) and *HBG1/2* promoters (sgHBG). The β-globin locus is flanked by two CTCF-associated insulator elements at HS5 and 3’HS1 ^36^. As important negative controls, we designed sgRNAs targeting DNA sequences 2∼4kb outside of the β-globin insulators (sgCTCF1 and sgCTCF2), outside of the β-globin locus-containing topologically associated domain (TAD) (sgTAD1 and sgTAD2), or at a different chromosome (sgCtrl; chr2:211,337,408-211,337,427) (Fig. 2e; Table S2). Upon stable co-expression of individual target-specific or control sgRNAs with enCRISPRi in K562 cells, we observed that sgHS2 resulted in more significant repression of all β-globin genes *HBE1, HBG1/2* and *HBB* (4.7 to 11.8-fold, *P* < 0.001 relative to sgGal4) compared to other LCR enhancers (Fig. 2f), consistent with the prominent role of the HS2 enhancer for LCR function ^35, 36^. Further, co-expression of all five sgRNAs targeting LCR enhancers (sgHS-all) did not further repress β-globin genes (Fig. 2e,f), suggesting that enCRISPRi was effective when targeted to a single enhancer by a single sgRNA. Notably, none of the control sgRNAs including two sgRNAs flanking β-globin insulators (sgCTCF1 and sgCTCF2) affected β-globin expression (Fig. 2e,f).

When targeted to gene-proximal promoters, dCas9-based epigenetic modulation may block TF binding and/or interfere with the formation of transcription initiation or elongation complexes ^17-19^. By contrast, enhancer repression requires the interference with the function of specific enhancer-regulating TFs, chromatin regulators and their combinatorial activities. We reasoned that targeting enCRISPRi to the proximity of enhancer center may achieve maximal effects compared to enhancer distal sequences. To this end, we compared the efficacies of enCRISPRi by designing sgRNAs targeting the DNA sequences at the DNase I hypersensitivity (DHS) peak summit at HS2 (sgHS2), or the sequences located 0.5kb or 2.5kb upstream or downstream of the HS2 enhancer (sgHS2-2.5k, sgHS2-0.5k, sgHS2+0.5k and sgHS2+2.5k; Fig. 2e), respectively. We achieved the most significant gene repression when targeting enCRISPRi to enhancer DHS peak summit (3.6 to 9.1-fold, *P* < 0.001 relative to sgGal4), and progressively decreased effects with increasing distances from enhancer center (Fig. 2f). These data emphasize the importance of targeting dCas9-based epigenetic editing complexes to the most accessible regions at enhancers for the maximal transcriptional perturbation.

Together, these results not only establish the improved epigenetic editing systems for enhancer perturbation, but also demonstrate that independent activators or repressors cooperate to modulate locus-specific gene transcription. It is important to note that, although the constituent components of enCRISPRa (p300 and VP64) and enCRISPRi (LSD1 and KRAB) have been tested for transcriptional modulation of promoter and/or enhancer activity by fusing to dCas9 individually ^18, 22, 41^, the combinatorial effects on local epigenetic landscapes and gene transcription have not been examined previously. Therefore, the enCRISPRa and enCRISPRi systems that we describe here represent the first attempt to combine both p300-VP64 and LSD1-KRAB in a single dCas9 complex for targeted modulation of enhancer function, respectively.

### Locus-Specific Epigenetic Editing by enCRISPRi

To determine the impact of enCRISPRi on epigenetic landscapes, we performed ChIP-seq analysis of dCas9, the enhancer-associated active histone marks (H3K4me1, H3K4me2 and H3K27ac), the repressive chromatin-associated H3K9me3, the hematopoietic lineage master TFs (GATA1 and TAL1), and CTCF. We also compared cells expressing non-targeting sgGal4 (control or C; Fig. 3), or sgHS2 with the single effector dCas9-KRAB (K), dCas9-LSD1 (L) or the dual effector enCRISPRi (LK or KL) (two replicate experiments for each ChIP-seq, total 80 independent ChIP-seq experiments; Figs. 3, S1, S2 and S3; Table S1).

**Figure 3.**
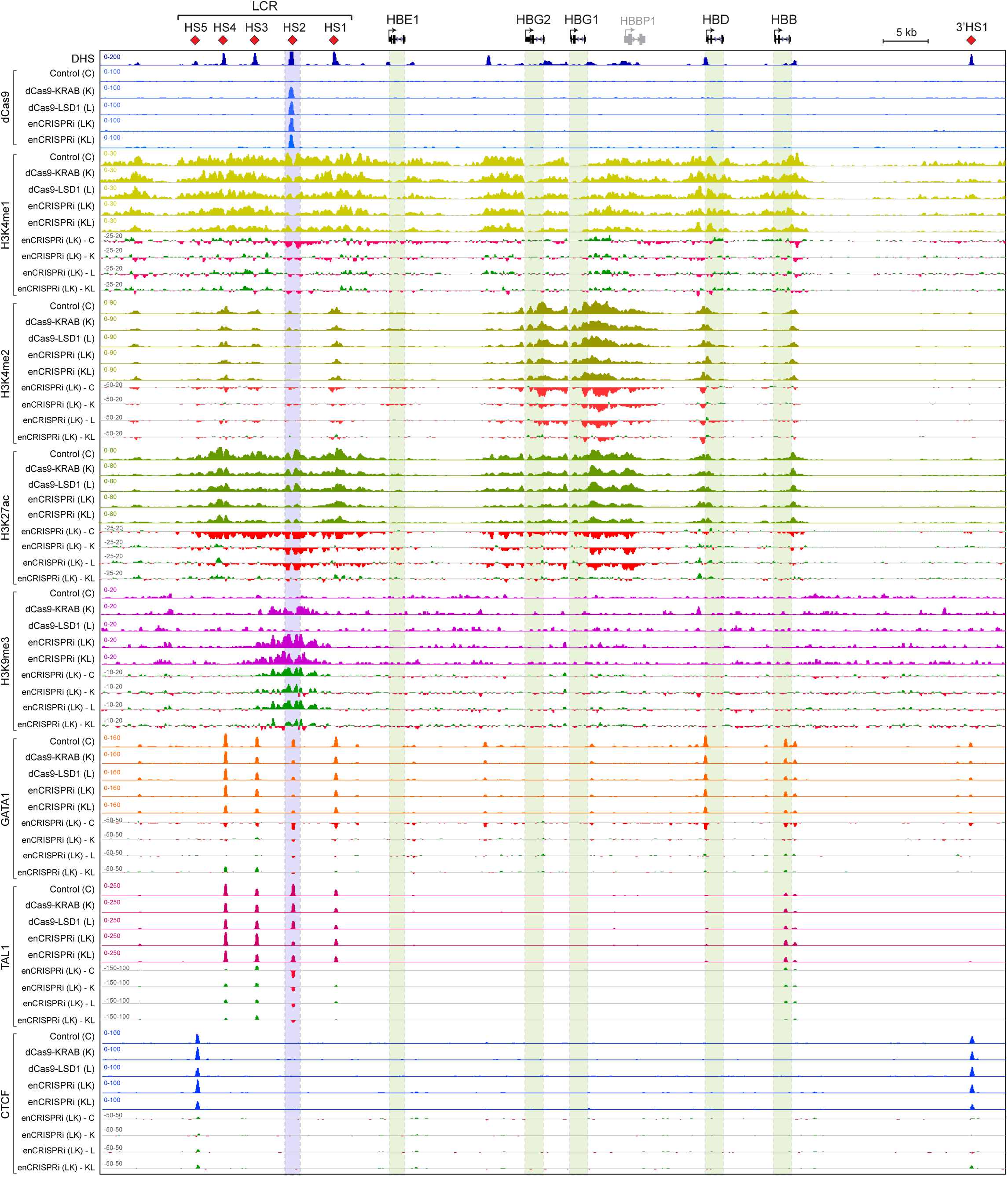
Locus-specific Epigenetic Reprogramming at the β-Globin Gene Cluster. Density maps are shown for ChIP-seq of dCas9, active histone marks (H3K4me1, H3K4me2, and H3K27ac), repressive H3K9me3, GATA1, TAL1, and CTCF at the β-globin cluster (chr11: 5,222,500-5,323,700; hg19) in K562 cells co-expressing non-targeting sgGal4 (control or C) or sgHS2 with dCas9-KRAB (K), dCas9-LSD1 (L) or enCRISPRi (LK and KL). Regions showing increased or decreased ChIP-seq signals in enCRISPRi (LK) relative to control, dCas9-KRAB, dCas9-LSD1 or enCRISPRi (KL) (enCRISPRi – C, enCRISPRi – K, enCRISPRi – L, or enCRISPRi – KL) are depicted in *green* and *red*, respectively. Blue bars denote the sgRNA-targeted HS2 enhancer. Green bars denote the β-globin genes. Independent replicate experiments are shown as rep1 and rep2 in Figs. S1 and S2, respectively.

Comparing to control (C), dCas9-LSD1 (L) or dCas9-KRAB (K), the dual repressor enCRISPRi LK and KL resulted in more apparent loss of enhancer-associated activating histone marks H3K4me1 and H3K27ac at the targeted HS2 enhancer (Figs. 3, S1 and S3a-d). Both enCRISPRi LK and KL significantly increased the levels of the repressive histone mark H3K9me3 at the targeted HS2 enhancer but not the β-globin promoters (Figs. 3, S1 and S3e). Of note, enCRISPRi also led to marked loss of H3K4me2 and H3K27ac at the β-globin gene-proximal promoters and gene bodies (Figs. 3 and S1), consistent with their transcriptional downregulation (Fig. 2b,c). These results suggest that enCRISPRi-mediated enhancer repression causes epigenetic changes at both targeted enhancers and associated gene promoters likely through enhancer-promoter interactions ^42^. It is also important to note that, while dCas9-KRAB (K) led to significantly increased H3K9me3 at the targeted HS2 enhancer, dCas9-LSD1 (L) had no effect on H3K9me3 but instead decreased H3K4me1/2 (Fig. 3). These results are consistent with the roles of KRAB in promoting the formation of H3K9me3-mediated heterochromatin ^30^ and LSD1 in the removal of H3K4me1/2 marks ^40^. Further, enCRISPRi (LK or KL) led to concurrent and more significant increases in H3K9me3 and decreases in H3K4me1/2 marks compared to dCas9-KRAB or dCas9-LSD1 alone (Figs. 3, S1 and S3b-e), illustrating the cooperative activity between two distinct repressor proteins. No significant enrichment of H3K27me3, the repressive histone marks catalyzed by Polycomb proteins ^43^, was detected at the β-globin gene cluster in K562 cells with or without dCas9-KRAB, dCas9-LSD1 or enCRISPRi (data not shown).

These data demonstrate that KRAB and LSD1 were capable of modulating epigenetic modifications by depositing H3K9me3 and removing H3K4me1/2, respectively, whereas the dual effector enCRISPRi led to more profound epigenetic changes, likely due to cooperation between distinct repressor domains. Finally, while dCas9-KRAB or dCas9-LSD1 alone slightly or modestly affected the binding of GATA1 and TAL1, the key hematopoietic TFs required for the HS2 enhancer function ^13, 35, 36^, enCRISPRi led to further loss of GATA1 and TAL1 binding at the targeted HS2 enhancer (Figs. 3, S2 and S3f,g). No significant effect on CTCF binding to the flanking insulators (HS5 and 3’HS1) was observed (Figs. 3, S2 and S3h), suggesting that enCRISPRi results in locus-wide epigenetic changes that are confined to the CTCF-bound insulated neighborhood.

Taken together, by focusing on the HS2 enhancer at the human β-globin gene cluster as a testbed, we provide evidence for the locus-specific epigenetic editing by enCRISPRi through KRAB-mediated H3K9me3 deposition and LSD1-mediated H3K4me1/2 removal. The dual effectors (KRAB and LSD1) act cooperatively to modulate locus-specific epigenetic modifications at the targeted enhancers and associated gene targets.

### Allele-Specific Activation of an Oncogenic Super-Enhancer by enCRISPRa

Having demonstrated the efficacy of the enhanced CRISPR perturbation systems *in vitro*, we examined whether we could modulate gene transcription and disease phenotypes *in vivo* by targeting disease-associated CREs. Recurrent mutations at an enhancer 8kb upstream of the *TAL1* proto-oncogene were discovered in T-cell acute lymphoblastic leukemia (T-ALL) cell lines and patients ^44^. In each case, the heterozygous somatic mutations are acquired through insertion of variable number of nucleotides at the *TAL1* enhancer sequences ^44^. In Jurkat T-ALL cells, a heterozygous 12bp insertion (GTTAGGAAACGG; Fig. 4a) introduces *de novo* binding motifs for the MYB proto-oncogene to initiate oncogenic super-enhancer (SE) formation ^44^. To establish the proof-of-principle for enCRISPRa in dissecting the *in vivo* role of *TAL1* oncogenic SE in T-ALL, we performed enCRISPRa-mediated activation of *TAL1* enhancer in Jurkat cells. We designed two independent sgRNAs that specifically target the mutant allele with protospacer-adjacent motifs (PAM) located within the 12bp insertion sequence (sgMut1 and sgMut2; Fig. 4a). As controls, we designed two sgRNAs targeting the wild-type enhancer sequences in the proximity of the 12bp insertion sequence (sgWT1 and sgWT2). By stable coexpression of enCRISPRa and individual sgRNAs targeting the *TAL1* oncogenic SE in Jurkat cells, we found that *TAL1* mRNA and protein were significantly induced by sgRNAs targeting WT or mutant alleles relative to the sgGal4 control (Fig. 4b,c). Since sgMut1 and sgMut2 specifically target the mutant enhancer sequences, these results suggest that allele-specific modulation of *TAL1* oncogenic SE allele achieves comparable efficacy on *TAL1* transcriptional activation as targeting both alleles.

**Figure 4.**
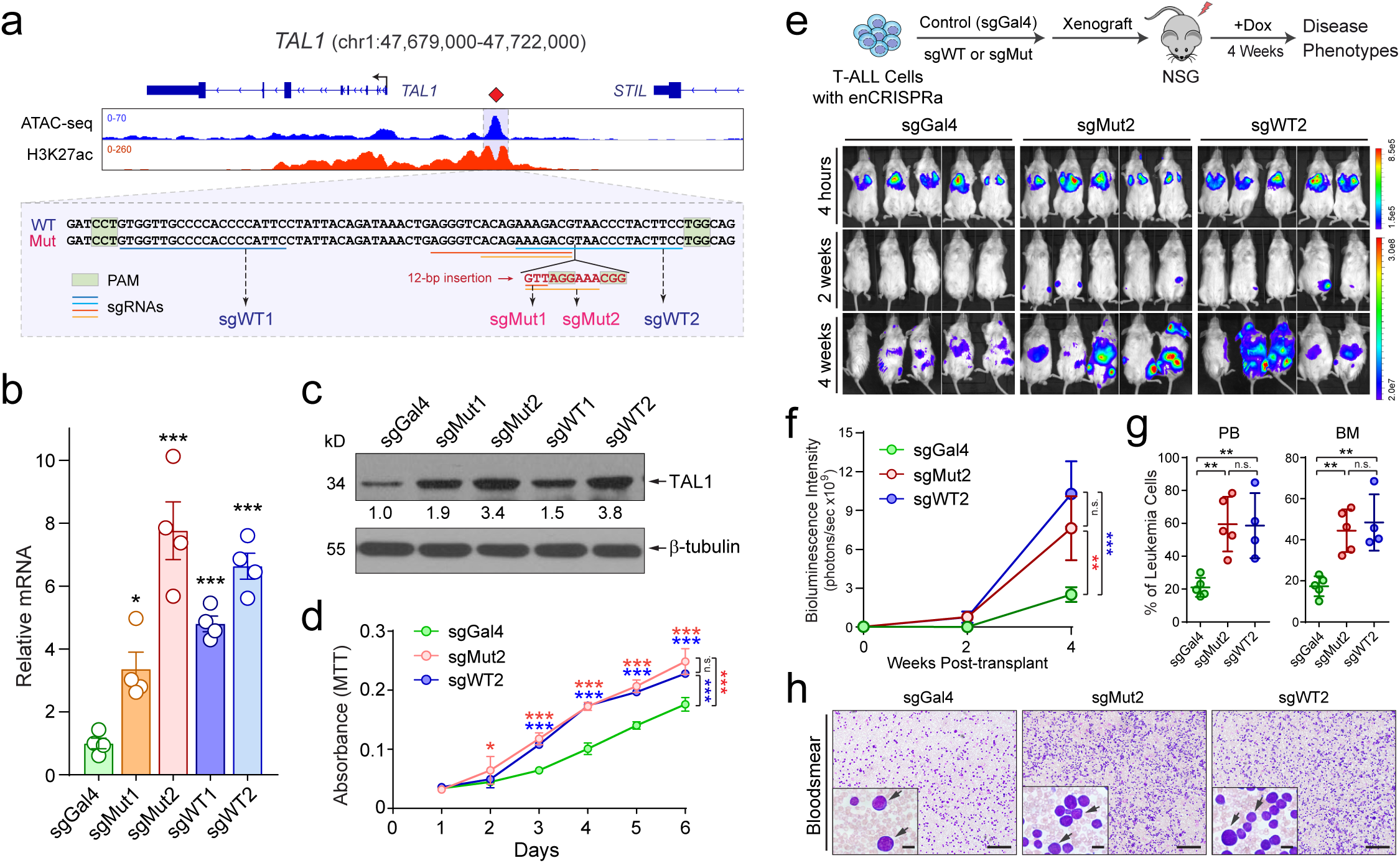
Activation of *TAL1* Oncogenic Super-Enhancer by enCRISPRa in Xenografts. **(a)** Density maps are shown for ATAC-seq and H3K27ac ChIP-seq at the *TAL1* locus (chr1:47,679,000-47,722,000; hg19) in Jurkat T-ALL cells. The annotated *TAL1* oncogenic SE is shown as blue shaded lines. WT or Mut enhancer sequences are shown in zoom-in view. The positions of sgRNAs targeting WT or Mut allele and corresponding PAM sequences are shown as colored lines and boxes, respectively. **(b)** Expression of *TAL1* mRNA in Jurkat cells upon enCRISPRa-mediated activation of *TAL1* oncogenic SE. Results are mean ± SEM (*N* = 4 independent experiments). **(c)** Expression of TAL1 protein by Western blot. The quantified TAL1 expression is shown. **(d)** Activation of *TAL1* oncogenic SE promoted T-ALL growth *in vitro*. Cell proliferation was determined by PrestoBlue cell viability assays and shown as relative absorbance (y-axis) after different days of culture (x-axis). **(e)** Activation of *TAL1* oncogenic SE promoted T-ALL growth in NSG mice xenografted with Jurkat cells transduced with control (sgGal4), sgMut2 or sgWT2, respectively. Bioluminescence intensity is shown for each mouse at 4 hours, 2- and 4-weeks post-transplantation. **(f)** Quantification of bioluminescence intensity is shown for the xenografted NSG mice. Results are mean ± SEM (*N* = 5 recipients per group). **(g)** Frequencies of leukemia cells in bone marrow (BM) and peripheral blood (PB) of xenografted NSG mice 4 weeks post-transplantation. Results are mean ± SEM (*N* = 5, 5, and 4 recipients for sgGal4, sgMut2 and sgWT2, respectively). **(h)** Representative bloodsmear images of NSG mice 4 weeks after xenotransplantation of control (sgGal4), sgMut2 or sgWT2-transduced Jurkat cells. The inset images indicate the zoom-in view. The representative leukemia cells are indicated by arrowheads. Scale bars, 200 µm and 20 µm for full and insert images, respectively. Results are mean ± SEM and analyzed by a one-way or two-way ANOVA for multiple comparisons. \**P* < 0.05, \*\**P* < 0.01, \*\*\**P* < 0.001, n.s. not significant.

Given the stronger effects on *TAL1* activation by sgMut2 and sgWT2 (Fig. 4a-c), we focused on these sgRNAs for the functional studies of enCRISPRa-mediated activation of *TAL1* oncogenic SE in T-ALL cells *in vitro* and *in vivo*. Importantly, enhanced *TAL1* expression by enCRISPRa led to significantly increased cell growth *in vitro* by PrestoBlue cell viability assay (see Methods; Fig. 4d), consistent with the oncogenic role of TAL1 in T-ALL cell proliferation ^44^. More importantly, upon xenotransplantation into immunodeficient NSG (NOD-*scid* IL2Rg^null^) mice, enCRISPRa-mediated *TAL1* enhancer activation led to greater tumor burden with significantly increased bioluminescence signals of the luciferase-expressing T-ALL cells in mice 4 weeks post-transplantation (Fig. 4e,f; see Methods). The mice transplanted with cells expressing *TAL1* SE-specific sgRNAs (sgMut2 and sgWT2) also displayed more severe leukemic phenotypes compared to the non-targeting sgGal4, resulting in increased infiltration of T-ALL leukemic cells in peripheral blood (PB) and bone marrow (BM) by flow cytometry and bloodsmear analyses (Fig. 4g,h; see Methods). These results not only establish the functional role of the *TAL1* oncogenic SE in promoting T-ALL development, but also demonstrate the efficacy of enCRISPRa for allele-specific activation of disease-associated enhancers *in situ* and *in vivo*.

### Generation of an Inducible Knock-In Mouse Model for *In Vivo* enCRISPRi

Systematic analysis of enhancer function *in vivo* remains a significant challenge. To explore the *in vivo* efficacy of enCRISPRi for enhancer perturbation, we engineered a new mouse model by site-specific knock-in (KI) of the dCas9-KRAB chimeric gene under the tetracycline-inducible promoter (TRE) into the *Col1a1* locus, which was previously used as a ‘safe harbor’ for robust transgene expression ^45, 46^, in KH2 embryonic stem cells (ESCs) (Fig. 5a; see Methods). KH2 ESCs also harbor the Rosa26-M2rtTA KI allele allowing expression of the rtTA-M2 trans-activator for doxycycline-inducible studies ^47^. After blastocyst injection of targeted ESCs and screening of germline transmitted offsprings, the founder mice were crossed to obtain the dCas9-KRAB::Rosa26-M2rtTA heterozygous or homozygous knock-in mice (named dCas9-KRAB KI hereafter) (Fig. 5b). The inducible dCas9-KRAB protein expression was confirmed in the targeted KH2 ESCs and bone marrow cells isolated from the dCas9-KRAB KI mice (Fig. 5c). To determine the effect of dCas9-KRAB expression on mouse hematopoietic development, we performed complete blood counts of peripheral blood (Fig. S4a) and flow cytometry of various mature hematopoietic cell types including erythroid (Ter119^+^), B-lymphoid (B220^+^), T-lymphoid (CD3^+^) and myeloid (Mac1+Gr1^+^) cells in bone marrow and spleen of wild-type (WT) or dCas9-KRAB KI mice with or without Dox treatment, respectively (Fig. S4b,c). Our results revealed no overt abnormalities on the frequency of mature hematopoietic cells before or after Dox-induced dCas9-KRAB expression (Fig. S4a-c). Furthermore, the cellularity and frequency of various hematopoietic stem/progenitor cell populations including HSC, MPP, LSK, CMP, GMP and MEP in bone marrow and spleen were comparable in WT and KI mice with or without Dox treatment (Fig. S5a,b; see Methods), indicating that the inducible dCas9-KRAB expression does not affect normal blood development.

**Figure 5.**
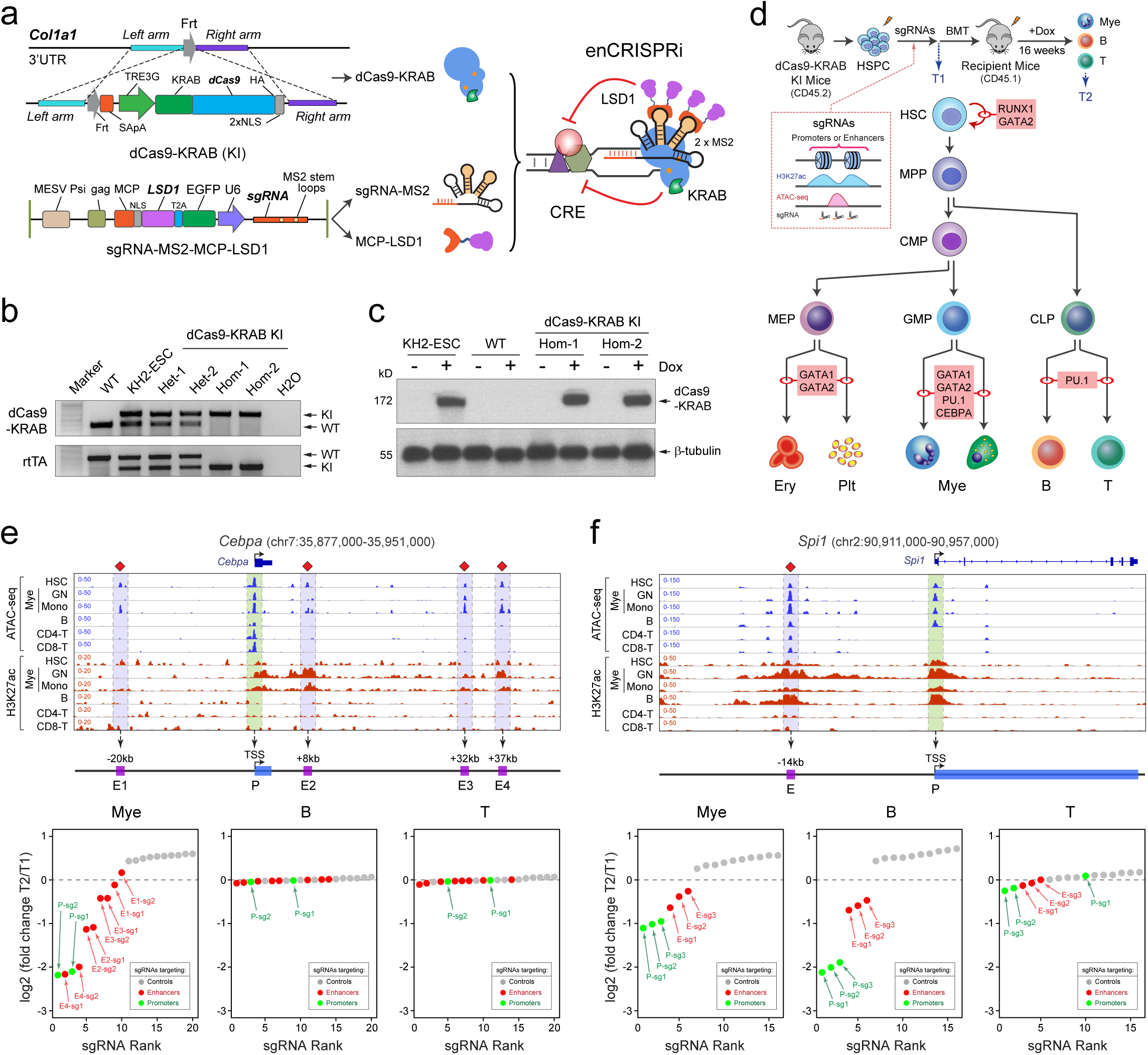
Locus-Specific *In Vivo* Enhancer Perturbation. **(a)** Schematic of site-specific KI of tetracycline-inducible dCas9-KRAB into the *Col1a1* locus. Co-expression of dCas9-KRAB, sgRNA and MCP-KRAB leads to assembly of enCRISPRi complex *in vivo*. **(b)** Validation of dCas9-KRAB and rtTA KI or WT alleles by genotyping PCR. C57BL/6 WT mouse and targeted KH2-ESC were used as controls. Two independent heterozygous (Het) and homozygous (Hom) KI mice were analyzed. **(c)** Dox-inducible expression of dCas9-KRAB fusion protein was confirmed by Western blot in the targeted KH2-ESC and two independent dCas9-KRAB KI mice. β-tubulin was analyzed as the loading control. **(d)** Schematic of *in vivo* perturbation of lineage-specific enhancers in dCas9-KRAB KI mice. **(e)** *In vivo* enCRISPRi perturbation of *Cebpa* CREs revealed lineage-specific requirement of *Cebpa* enhancers during hematopoiesis. Waterfall plots are shown for target-specific sgRNAs (*green* and *red* dots) and non-targeting control sgRNAs (*grey* dots) by the mean normalized log2 fold changes in myeloid, T or B cells 16-weeks post-BMT (T2) relative to pooled sgRNA-transduced HSPCs (T1) from two independent replicate screens (*N* = 3 recipient mice per screen). Density maps are shown for ATAC-seq and H3K27ac ChIP-seq at the *Cebpa* locus (chr7:35,877,000-35,951,000; mm9) in bone marrow HSC, granulocytes (GN), monocytes (Mono), B, CD4+ and CD8+ T cells, respectively. The annotated *Cebpa* promoter (P) and enhancers (E1 to E4) are indicated by *green* and *blue* shaded lines. Results from independent replicate screens and statistical analyses are shown in Fig. S6a,b. **(f)** *In vivo* enCRISPRi perturbation of *Spi1* CREs during hematopoiesis. Density maps are shown for ATAC-seq and H3K27ac ChIP-seq at the *Spi1* locus (chr2:90,911,000-90,957,000; mm9) in bone marrow HSC, GN, Mono, B, CD4+ and CD8+ T cells, respectively. The annotated *Spi1* promoter (P) and enhancer (E) are indicated by *green* and *blue* shaded lines. Results from independent replicate screens and statistical analyses are shown in Fig. S6c,d.

Hematopoiesis serves as a paradigm for understanding stem cell differentiation controlled by lineage-specifying TFs ^48^. The self-renewing hematopoietic stem cells (HSCs) give rise to all mature blood cell lineages through a hierarchy of progenitors during lineage specification. In the bone marrow transplantation (BMT) setting, HSCs are capable of reconstituting the entire blood system of a recipient, whereas the short-lived progenitors do not. HSC self-renewal and/or lineage determination are controlled by a small number of TFs, many of which function in a highly lineage-specific manner and are regulated by tissue-specific and/or developmentally regulated enhancers ^48^. We reasoned that by epigenetic modulation of lineage-specific enhancers in HSCs followed by BMT, we could assay the HSC-derived mature cell lineages as the ‘readout’ for the functional impact of enhancer perturbations during HSC lineage differentiation *in vivo*.

### *In Vivo* Functional Interrogation of Lineage-Specific Enhancers

To this end, we devised an *in vivo* enhancer perturbation assay by combining the dCas9-KRAB KI mice with sgRNA-MS2 and MCP-LSD1 to assemble the enCRISPRi complex *in vivo* (Fig. 5a). We then determined the functional role of lineage-specific enhancers associated with major hematopoietic TFs by *in vivo* enCRISPRi (Fig. 5d). We focused on five TFs including *Cebpa* ^49^, *Spi1* (or PU.1) ^50^, *Gata1* ^51^, *Gata2* ^52, 53^ and *Runx1* ^54, 55^ that play critical roles in HSC function and/or lineage differentiation ^48^. Each gene contains one or multiple annotated enhancers based on chromatin accessibility by ATAC-seq and enhancer-associated H3K27ac by ChIP-seq ^56^ (Figs. 5e,f, 6c, and S6-S8). We designed 2 or 3 sgRNAs for each enhancer, 2 or 3 sgRNAs for each gene promoter as positive controls, and 10 non-targeting sgRNAs as negative controls (Table S2). The target-specific and control sgRNAs were pooled for each gene for locus-specific enhancer perturbation (total 5 pools with 16 to 20 sgRNAs in each pool). CD45.2^+^ BM hematopoietic stem/progenitor cells (HSPCs) from dCas9-KRAB KI mice were isolated and retrovirally transduced with pooled sgRNAs at MOI ≤ 0.5 to ensure that each cell contained no more than one sgRNA (see Methods). The transduced cells were selected and transplanted into CD45.1^+^ lethally irradiated recipient mice, followed by Dox administration to induce dCas9-KRAB expression for 16 weeks. By this time, all donor-derived hematopoietic cells (CD45.2^+^) were differentiated from repopulating donor HSCs instead of short-lived progenitors. We collected cells before BMT (T1) as the baseline control and donor-derived mature myeloid (Gr1^+^Mac1^+^), B (B220^+^) and T (CD3^+^) lymphoid cells in the peripheral blood of recipient mice 16 weeks post-BMT (T2). We did not analyze erythroid and megakaryocytic lineages because mature erythroblasts and platelets do not contain nuclei. Genomic DNA from T1 and T2 cells were isolated to amplify the sgRNA sequences, followed by high-throughput amplicon sequencing (see Methods). If enCRISPRi-mediated repression of an enhancer or promoter impaired its function and target gene expression, the corresponding sgRNAs would be depleted (or enriched) in T2 relative to T1 cells (Fig. 5d).

**Figure 6.**
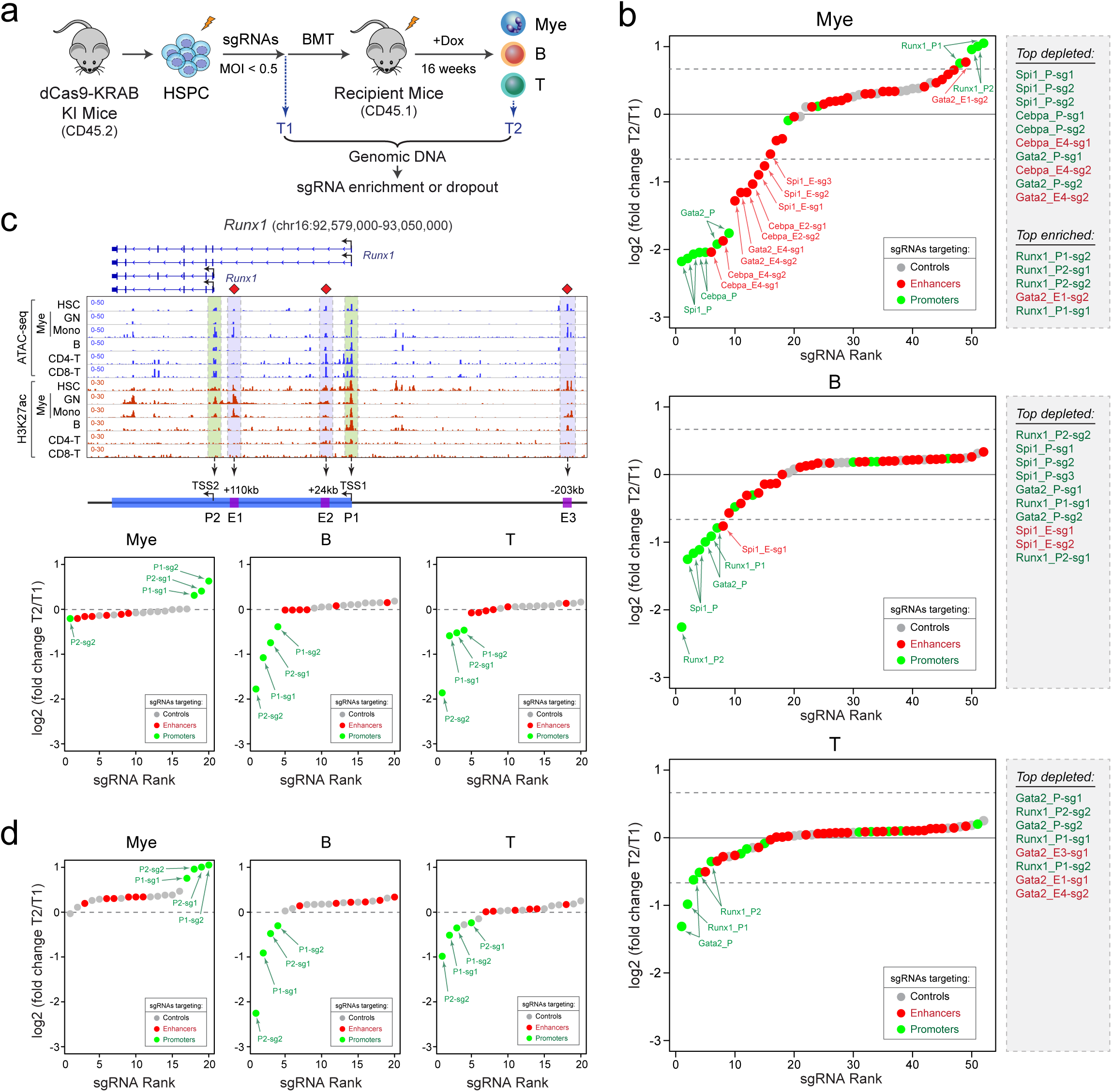
Multiplexed Perturbation of Developmental Enhancers During Hematopoiesis. **(a)** Schematic of *in vivo* multiplexed perturbation of developmentally regulated enhancers in dCas9-KRAB KI mice. **(b)** *In vivo* perturbation of annotated CREs for five key hematopoietic TFs revealed the functional requirement of lineage-specific enhancers for HSC differentiation to one or multiple hematopoietic lineages. Waterfall plots are shown for target-specific sgRNAs (*green* and *red* dots) and non-targeting control sgRNAs (*grey* dots) by the mean normalized log2 fold changes in myeloid, T or B cells 16-weeks post-BMT (T2) relative to pooled sgRNA-transduced HSPCs (T1) from two independent replicate screens (*N* = 15 recipient mice per replicate screen). The annotated *Runx1* promoters (P1 and P2) and enhancers (E1 to E3) are indicated by *green* and *blue* shaded lines. Results from independent replicate screens and statistical analyses are shown in Fig. S10. **(c)** *In vivo* enCRISPRi of *Runx1* CREs during hematopoiesis by single locus-based perturbation. Density maps are shown for ATAC-seq and H3K27ac ChIP-seq at the *Runx1* locus (chr16:92,579,000-93,050,000; mm9) in bone marrow HSC, GN, Mono, B, CD4+ and CD8+ T cells, respectively. Results from independent replicate screens and statistical analyses are shown in Fig. S8a,b. **(d)** *In vivo* enCRISPRi of *Runx1* CREs during hematopoiesis by multiplexed perturbation. Results from independent replicate screens and statistical analyses are shown in Fig. S8c,d.

We performed five independent locus-specific enhancer perturbation screens, each with two independent replicate experiments, for *Cebpa, Spi1, Gata1, Gata2* and *Runx1*, respectively (Figs. 5e,f, 6c, and S6-S8). Deletion of *Cebpa* or its downstream enhancer (E4, +37kb from TSS; Fig. 5e) led to defective myeloid-lineage priming and differentiation without affecting lymphopoiesis ^49, 57^. Consistent with these findings, sgRNAs for E4 enhancer were the top depleted sgRNAs in myeloid cells (ranked #2 and #4) next to the positive control sgRNAs for the *Cebpa* promoter (ranked #1 and #3; Fig. 5e) in independent screen experiments (Fig. S6a,b). More importantly, our results revealed that *Cebpa* E2 (+8kb) enhancers was also required for *in vivo* myelopoiesis, whereas E1 (−20kb) and E3 (+32kb) were dispensable (Figs. 5e and S6a,b). None of the *Cebpa* enhancers was required for B or T cell development considering both the fold changes of sgRNA abundance and the significance of sgRNA depletion between T1 and T2 (Figs. 5e and S6a,b; see Methods). For *Spi1* locus, we observed significant depletion of all 3 independent sgRNAs for its −14kb enhancer in myeloid and B but not T cells (Figs. 5f and S6c,d). These results are consistent with the role of Spi1/PU.1 for normal myeloid and B cell development ^50, 58^, and validate the *in vivo* requirement of its upstream enhancer in regulating myelopoiesis and B lymphopoiesis ^59^.

With regards to *Gata1*, while sgRNAs for *Gata1* promoters were modestly depleted in myeloid cells consistent with the role of GATA1 in eosinophil and mast cell function ^48, 60^, none of the sgRNAs for two *Gata1* enhancers was depleted in any of the three lineages (Fig. S7a,b). Interestingly, *Gata1* enhancers displayed chromatin accessibility and H3K27ac enrichment in HSCs but not mature myeloid, B and T cells, suggesting that the HSC-specific *Gata1* enhancers are not required for HSC differentiation to mature myeloid or lymphoid lineages. For *Gata2*, we observed that an intronic enhancer (+9.5kb, E4) was required for *in vivo* myelopoiesis, whereas the other three gene-distal enhancers were minimally required for any of the three lineages (Fig. S7c,d). It is important to note that the *in vivo* functional impacts of enhancer perturbation largely correlated with the levels of target gene repression by enCRISPRi-mediated targeting of individual enhancers or promoters in HSPCs (Fig. S9). The lineage-restricted impacts on HSC differentiation also correlated with the presence of enhancer-associated chromatin features (ATAC-seq and H3K27ac) in different lineages (Figs. 5e,f, 6c, and S6-S8), highlighting the context-specific requirements of developmental enhancers in regulating gene expression and lineage differentiation.

### Multiplexed Perturbations of Enhancer Function During Hematopoiesis

Locus-specific perturbation screens provided information for the ‘essentiality’ of each enhancer within its local context (Figs. 5, S6 and S7), but did not reveal the relative importance of enhancers across multiple loci during hematopoiesis. To address this question, we pooled the target-specific and control sgRNAs for all five hematopoietic TFs, and performed multiplexed enhancer perturbation screens (Fig. 6a). We observed that the top depleted enhancer sgRNAs in myeloid cells included *Cebpa* +37kb (E4) and +8kb (E2) enhancers, *Gata2* +9.5kb (E4) and *Spi1* −14kb enhancers when considering the fold changes of sgRNA abundance and the significance of sgRNA depletion (Figs. 6b and S10). The sgRNAs for *Spi1* and *Gata2* E1, E3 and E4 enhancers were modestly or slightly depleted in B and T cells, respectively (Figs. 6b and S10). These results illustrate the disparate requirements for distinct enhancers during hematopoiesis, with some enhancers broadly required for multiple lineages (e.g. *Gata2* or *Spi1* enhancers) while others uniquely required for one specific lineage (e.g. *Cebpa* enhancers). Our results also demonstrate that the *in vivo* enhancer functions cannot be solely predicted based on enhancer-associated epigenetic features. For example, while all four *Cebpa* enhancers share similar chromatin accessibility and H3K27ac enrichment in HSCs and/or myeloid cells, only +8kb E2 and +37kb E4 enhancers are indispensable for myeloid development (Figs. 5e and 6b).

By locus-specific enhancer perturbation, we found that sgRNAs for *Runx1* promoters (P1 and P2) were significantly depleted in B and T cells but enriched in myeloid cells, whereas none of sgRNAs for the annotated *Runx1* enhancers showed differential enrichment (Figs. 6c and S8a,b). Similar results were replicated in multi-loci multiplexed perturbation (Figs. 6d and S8c,d). These findings demonstrate that enCRISPRi-mediated repression of *Runx1* promoters impaired HSC differentiation to B and T lineages but promoted myeloid differentiation; however, enCRISPRi-mediated enhancer repression was ineffectual, supporting the non-essential roles of *Runx1* enhancers in myelopoiesis or lymphopoiesis. More importantly, the opposing phenotypes of repressing *Runx1* by enCRISPRi in myeloid vs lymphoid lineages are consistent with the differential roles of *Runx1* in hematopoiesis ^61^. Hematopoietic cell-specific *Runx1* KO in mice led to the development of myeloproliferative phenotypes characterized by defective B- and T-cell maturation, and increased myelopoiesis ^61^. Our results faithfully recapitulated the phenotypic manifestations of *Runx1* deficiency, illustrating the utility of enCRISPRi-based epigenetic editing in functional analysis of *cis*-regulatory elements during *in vivo* development. Hence, the exemplified applications of enhancer perturbations *in vitro* (Figs. 1-3), in xenografts (Fig. 4) and *in vivo* (Figs. 5, 6) in multiple cell models establish the enhanced CRISPR epigenetic editing systems as general tools for functional interrogation of non-coding regulatory genome in development and disease.

## DISCUSSION

### Targeted Epigenetic Editing *In Situ* by Dual Effector dCas9 Complexes

Here we describe the enhanced CRISPR epigenetic editing systems for locus-specific activation or repression of transcriptional enhancers or other CREs in native chromatin. Using epigenetic writer proteins that specifically modulate histone modifications associated with active enhancers, the new systems enable highly efficient and multiplexed analysis of enhancer function in human and mouse cells *in vitro*, in xenografts and *in vivo*. Our results also demonstrate that independent repressors (LSD1 and KRAB in enCRISPRi) or activators (p300 and VP64 in enCRISPRa) cooperate to modulate gene transcription by remodeling epigenetic landscapes at the targeted genomic loci. KRAB is associated with heterochromatin formation ^30^, whereas LSD1 removes enhancer-associated H3K4me1/2 ^40^. The combined effects on H3K9me3 deposition and H3K4me1/2 removal exceeded individual effectors when targeted to the HS2 enhancer (Figs. 3, S1 and S3), suggesting that different epigenetic effectors act cooperatively and/or synergistically for maximal enhancer perturbations. Moreover, the development of enCRISPRi and enCRISPRa systems permits parallel analyses of the same loci by both loss- and gain-of-function approaches, thus facilitating the identification of CREs that are necessary and/or sufficient for target gene expression. It is important to note that, while our studies focus on the applications of enCRISPRi and enCRISPRa for modulating enhancer activities, the dual-effector systems also work effectively on modulating gene expression when targeted to gene-proximal promoters *in vitro* and *in vivo* (Figs. 1b, 5e,f, 6b-d, S6-S8 and S10).

### Considerations for Enhancer Perturbations by CRISPR Epigenetic Editing

We found that a single sgRNA targeting a single enhancer was sufficient for gene repression; however, the position of sgRNAs can significantly affect the effectiveness of CRISPR epigenetic editing as shown in the case of HS2 enhancer (Fig. 2e,f). Given that we positioned sgRNAs based on the distances to the DHS or ATAC-seq peak summits at enhancers, these results indicate that efficient enhancer perturbations require maximal interference with the chromatin binding of enhancer-regulating TFs. Consistent with this idea, the dual effector enCRISPRi led to more significant disruption of GATA1 and TAL1 binding than single effector dCas9-KRAB or dCas9-LSD1 at HS2 enhancer (Fig. 3). These findings are distinct from CRISPRi in promoter repression, in which targeting sgRNAs to promoter downstream sequences (50∼100bp) was most effective in gene repression likely due to the interference with transcription initiation and/or elongation complexes ^19^. In future studies, high-resolution tiling screens of critical TF binding site(s) at enhancer sequences may elicit the functional roles of individual TF binding sites in enhancer regulation. Such studies will not only provide insights into the regulatory mechanisms for enhancer function, but also identify selective ‘vulnerabilities’ of enhancers that may be employed to precisely control gene expression.

### *In Vivo* Interrogation of Enhancer Functions in Development and Disease

By *in vivo* multiplexed perturbation screens, we identified several candidate enhancers required for lineage differentiation of HSCs (Figs. 5, 6 and S6-S10). Although the lineage-specific essentiality largely correlated with the presence of enhancer-associated chromatin accessibility and H3K27ac, the functional roles of individual enhancers cannot be reliably predicted based on only epigenetic features. For instances, while the annotated *Cebpa* enhancers share similar chromatin features, only +8kb E2 and +37kb E4 enhancers are indispensable for *Cebpa* expression and myeloid development (Figs. 5e, 6b and S6). Similarly, while several annotated *Runx1* enhancers harbor enhancer-associated chromatin features in HSCs and/or differentiated lineages, none of them was required for *Runx1* expression or function in HSC differentiation to myeloid and lymphoid cells *in vivo* (Figs. 6 and S8). These results highlight the importance of analyzing enhancer function *in situ* by loss- and gain-of-function assays, such as the enCRISPRi and enCRISPRa epigenetic editing systems described here, ideally during *in vivo* development. These assays require the analysis of cellular phenotypes such as stem cell differentiation, cell proliferation, viability, and/or response to stimuli as readouts to quantify the functional impact.

Therefore, the development of dCas9-KRAB KI mouse model provides opportunities for *in vivo* functional interrogation of enhancers and other CREs in lineage differentiation of tissue stem cells. Multiplexed analysis of many genes or loci allows efficient functional screens to prioritize relevant CREs for in-depth characterization. In future studies, the inducible dCas9-KRAB KI mouse model combined with various disease models will provide new insights into the roles of non-coding CREs in disease pathogenesis *in vivo*. In addition, the improved CRISPR epigenetic editing systems should accelerate functional follow-up studies of disease or trait-associated genetic variants and cancer-associated somatic alterations, many of which reside in non-coding CREs including enhancers. Finally, the enhanced CRISPR epigenetic editing may suggest potential therapeutic strategies by generating targeted epigenetic modifications to alter the expression of desired genes. In conclusion, we have developed dual activator or repressor-containing CRISPR perturbation systems for functional analysis of non-coding regulatory elements. Our studies not only identify candidate lineage-specific enhancers required for hematopoiesis, but also establish a widely applicable platform for unbiased analysis of non-coding regulatory genome which can be extended to other cell types and human diseases.

## AUTHOR CONTRIBUTIONS

Conceptualization, K.L., Y.L. and J.X.; Methodology, K.L., Y.L., H.C., Y.Z., Z.G., X.L., A.Y., P.K., K.E.D., M.N. and J.X.; Investigation, K.L., X.L., H.C., P.K. and J.X.; Writing – Original Draft, K.L., Y.L. and J.X.; Writing – Review & Editing, J.X.; Funding Acquisition, J.X.; Supervision, J.X.

## ACKNOWLEDGMENTS

We thank the BioHPC computational infrastructure at UTSW for assistance, Lin Li and Le Qi for technical support, and Luke A. Gilbert, Stanley S. Qi, and Jonathan S. Weissman at UCSF for providing the CRISPRi constructs. K.L. and Y.L. were supported by the Cancer Prevention and Research Institute of Texas (CPRIT) training grant (RP160157). X.L. was supported by the American Heart Association postdoctoral fellowship (18POST34060219). J.X. is a Scholar of The Leukemia & Lymphoma Society. This work was supported by the NIH grants R01DK111430 and R01CA230631, the CPRIT grants (RR140025, RP180504, RP180826 and RP190417), and the Welch Foundation grant I-1942 (to J.X.).

## METHODS

### Cells and Cell Culture

Human K562 and Jurkat cells were obtained from ATCC and cultured RPMI1640 medium containing 10% fetal bovine serum (FBS), 1% penicillin/streptomycin (P/S). Human HEK293T cells were obtained from ATCC and cultured in DMEM medium containing 10% FBS and 1% P/S. KH2 ESCs were cultured in DMEM medium containing 10% ES-certified FBS (GemCell™, Cat# 100-500), 1% P/S, 2mM L-Glutamine (Gibco), 0.1mM MEM non-essential amino acids solution (Gibco), 1mM sodium pyruvate (Gibco), 0.1mM β-Mercaptoethanol and 1000U/ml recombinant mouse LIF (ESGRO, Cat# ESG1107). All cultures were incubated at 37°C in 5% CO2. No cell line used in this study was found in the database of commonly misidentified cell lines that is maintained by ICLAC and NCBI BioSample. All cell lines were tested for mycoplasma contamination.

### Plasmids

To generate the inducible dCas9-p300 expression vector, the p300 HAT core domain (p300 core) was PCR amplified from the pcDNA-dCas9-p300-Core vector (Addgene, Plasmid #61357) and cloned into MluI/BstXI digested pHR-TRE3G-KRAB-dCas9-P2A-mCherry backbone, which was a gift from Luke A. Gilbert ^23^. To generate the inducible dCas9-LSD1 expression vector, LSD1 open-reading frame (ORF) was amplified and cloned into the pHR-TRE3G-KRAB-dCas9-P2A-mCherry to replace the KRAB domain. To generate the enCRISPRa sgRNA vector, the MCP-VP64-IRES-mCherry cassette was PCR amplified from the pJZC34 vector (Addgene, plasmid # 62331) and cloned into BsrGI/EcoRI digested lenti-sgRNA (MS2)-zeo backbone (Addgene, plasmid # 61427). Then the mCherry cassette was replaced by Puro or EGFP by In-Fusion^®^ HD Cloning Kit (Clontech). To generate the enCRISPRi-LK sgRNA vector, the KRAB sequence was PCR amplified from the pLV hU6-sgRNA hUbC-dCas9-KRAB-T2A-Puro vector (Addgene, plasmid #71236) and cloned into the enCRISPRa sgRNA vector to replace VP64. To generate enCRISPRi-KL sgRNA vector, MCP-LSD1 was amplified and cloned into pMLS-NRAS-T2A-GFP-polyA-U6 to replace NRAS.

### Design and Cloning of sgRNAs

sgRNAs were designed to minimize off-targets based on publicly available filtering tools (http://crispr.genome-engineering.org/crispr/). Briefly, oligonucleotides were annealed in the following reaction: 10μM guide sequence oligo, 10μM reverse complement oligo, T4 ligation buffer (1X), and 5U of T4 polynucleotide kinase (NEB) with the cycling parameters of 37°C for 30 min; 95°C for 5 min and then ramp down to 25°C at 5°C/min. The annealed oligos were cloned into the sgRNA vectors using a Golden Gate Assembly strategy including: 100ng of circular sgRNA vector plasmid, 0.2μM annealed oligos, buffer 2.1 (1X) (NEB), 20U of BbsI restriction enzyme, 0.2mM ATP, 0.1mg/ml BSA, and 750U of T4 DNA ligase (NEB) with the cycling parameters of 20 cycles at 37°C for 5 min, 20°C for 5 min; followed by 80°C incubation for 20min. Insertion of sgRNA was validated by Sanger sequencing. Lentiviruses containing sgRNAs were packaged in HEK293T cells. Briefly, 2µg of pΔ8.9, 1µg of VSV-G and 5µg sgRNA vectors were co-transfected into HEK293T cells seeded in 10cm petri dish. Lentiviruses were harvested from the supernatant 48∼72 h post-transfection. Dox-Inducible enCRISPRi or enCRISPRa cell lines were then transduced with sgRNA-expressing lentiviruses in 6-well plates. To maximize sgRNA expression, top 1-5% of GFP-positive cells were FACS sorted 48 h post-infection. The sequences for all sgRNAs are listed in Table S2.

### Generation of Tetracycline-Inducible dCas9-KRAB Knock-In Mouse Model

Site-specific knock-in (KI) of tetracycline-inducible dCas9-KRAB transgene was generated by flippase (FLPe)-mediated recombination as previously described ^35, 62^. KH2 mouse embryonic stem cells (ESCs) harboring the M2rtTA tetracycline-responsive trans-activator in *Rosa26* locus and an engineered *Col1a1* locus with an frt site and ATG-less hygromycin resistance gene were used ^62^. A targeting construct pBS3.1-TRE-dCas9-KRAB containing the PGK promoter, an frt site, a tetracycline-inducible minimal CMV promoter, the dCas9-KRAB transgene, and an ATG initiation codon was co-electroporated with the pCAGGS-FLPe-puro into KH2 ESCs at 500V and 25 µF using a Gene Pulser II (Bio-Rad). Cells were selected with hygromycin (140 µg/ml), and positive clones were expanded and analyzed by genotyping PCR. Targeted ESC clones were injected to embryonic day 3.5 mouse blastocysts to obtain the founder mice. Chimeric founder mice were bred with C57BL/6 mice, and offsprings with germline transmission were genotyped using primers in Table S2 and intercrossed to generate dCas9-KRAB heterozygous or homozygous KI mice. All mouse experiments were performed under protocols approved by the Institutional Animal Care and Use Committee of UT Southwestern Medical Center (UTSW).

### Xenograft Experiments

Luciferase cassette was amplified and cloned into pLVX-Puro vector (Clontech, Catalog No. 632164). Lentivirus was produced to transduce Jurkat cells co-expressing enCRISPRa with control sgGal4, sgWT2 or sgMut2, respectively. Puromycin selection (1 µg/ml) was performed 3 days after infection. Six to eight weeks old female NOD-SCID (NSG) mice were sub-lethally irradiated (2.5 Gy) half day before the transplantation. Cells (1×10^6^/mice) were resuspended in PBS (200µl/mice) and intravenously transplanted. Transplanted mice underwent *in vivo* bioluminescence imaging at various time points to evaluate tumor growth. Briefly, following intraperitoneal injection of 150mg/kg D-luciferin (Gold Biotechnology), mice were imaged, and bioluminescence intensity was quantitated using Living Image 3.2 acquisition and analysis software (Caliper Life Sciences). Total flux values were determined by drawing regions of interest (ROI) of identical size over each mouse and were presented in photons (p)/second (sec). Four weeks after transplantation, the peripheral blood, bone marrow and spleen were assessed for engraftment by flow cytometry. Bloodsmear was performed and stained with May-Grunwald-Giemsa as previously described ^63^. The blue stained cells indicated the circulating leukemia cells.

### Generation of Dox-Inducible enCRISPRi or enCRISPRa Cell Lines

To generate inducible enCRISPRi and enCRISPRa stable cell lines, the target cells were transduced with lentivirus expressing dCas9-KRAB, dCas9-LSD1 or dCas9-p300 and rtTA. Doxycycline was added following infection and flow cytometry was used to sort cells that expressed mCherry and BFP. These cells were then grown in the absence of doxycycline until mCherry fluorescence returned to uninduced levels. Transient transfections were performed in 24-well plates using 500ng of dCas9 expression vector and 250ng of equimolar pooled or individual sgRNA expression vectors mixed with FuGENE^®^ 6 (Promega) following manufacturer’s instructions.

### Cell Growth Assays

Cell proliferation was determined using the PrestoBlue Cell Viability Reagent (Invitrogen). 5,000 cells/well were seeded in triplicate into 96-well plates. After various days of culture, 10μl of PrestoBlue reagents were added to wells with cells or medium (blank), relative absorption values at 570 and 600 nm were read after 1 hour incubation at 37°C.

### RNA Isolation and qRT-PCR Analysis

RNA was isolated using RNeasy Plus Mini Kit (Qiagen) following manufacturer’s protocols. For qRT-PCR, RNA was reverse-transcribed using iScript cDNA Synthesis Kit (Bio-Rad) following manufacturer’s protocols. Quantitative RT-PCR (qRT-PCR) was performed in duplicate with the iQ SYBR Green Supermix (Bio-Rad) using CFX384 Touch Real-Time PCR Detection System (Bio-Rad). PCR amplification parameters were 95°C (3 min) and 45 cycles of 95°C (15 sec), 60°C (30 sec), and 72°C (30 sec). Primer sequences are listed in Table S2.

### Western Blot Analysis

Western blot was performed as described ^64^ using the following antibodies: TAL1 (Santa Cruz Biotechnology, sc-12984), β-tubulin (Cell Signaling, 2128), HA (Santa Cruz Biotechnology, sc-805), and Cas9 (Abcam, ab191468). All antibodies were used at 1:1,000 dilutions. Densitometry quantification was performed using ImageJ software.

### Phenotypic Analysis of Hematopoiesis

Blood was collected via the retro-orbital plexus and complete blood counts (CBC) were performed on a HEMAVET HV950 (Drew Scientific) according to the manufacturer’s protocol. Cytospin preparations were stained with May-Grunwald-Giemsa as described previously ^65^. BM cells were obtained by crushing femurs, tibias, vertebrae and pelvic bones with a mortar in Ca^2+^ and Mg^2+^-free Hank’s buffered salt solution (HBSS, Gibco) supplemented with 2% heat-inactivated bovine serum (HIBS, Gibco). Spleens were dissociated by crushing followed by trituration. All BM and spleen cell suspensions were filtered through 70 μm cell strainers, followed by cell counting using a Vi-CELL cell viability analyser (Beckman Coulter). For flow cytometric analysis, cells were incubated with combinations of fluorophore-conjugated antibodies. Lineage markers for HSCs and progenitors were CD2, CD3, CD5, CD8, B220, Gr1 and Ter119. Antibody staining was performed at 4 °C for 30 min or on ice for 90 min. Biotinylated antibodies were visualized by incubation with PE/Cy7-conjugated streptavidin at 4°C for 30 min. DAPI (4,6-diamidino-2-phenylindole; 2 μg/ml in PBS) was used to exclude dead cells. The hematopoietic stem/progenitor cell populations in mouse bone marrow were analyzed as previously described ^63^, including HSC (Lin^-^Sca1^+^Kit^+^CD150^+^CD48^-^), MPP (Lin^-^ Sca1^+^Kit^+^CD150^-^CD48^-^), LSK (Lin^-^Sca1^+^Kit^+^), CMP (Lin^-^Sca1^-^Kit^+^CD34^+^CD16/32^-^), GMP (Lin^-^ Sca1^-^Kit^+^CD34^+^CD16/32^+^), and MEP (Lin^-^Sca1^-^Kit^+^CD34^-^CD16/32^-^). Analysis or sorting was performed using a FACSAria or FACSCanto flow cytometer (BD Biosciences). Data were analyzed using FACSDiva (BD Biosciences).

The following antibodies were used for flow cytometry: PerCP/Cy5.5 anti-mouse B220 (Biolegend, Cat# 103236), PE anti-mouse CD150 (Biolegend, Cat# 115903), BV510 anti-mouse CD16/32 (Biolegend, Cat# 101333), Alexa Fluor 700 anti-mouse CD3 (Biolegend, Cat# 100216), Biotin anti-mouse CD34 (eBioscience, Cat# 13-0341-85), PE anti-mouse CD43 (eBioscience, Cat# 12-0431-83), Alexa Fluor 700 anti-mouse CD48 (Biolegend, Cat# 103426), FITC anti-mouse CD71 (BD Biosciences, Cat# 553266), APC anti-mouse c-Kit (Biolegend, Cat# 105811), PE/Cy7 anti-mouse Gr1 (Biolegend, Cat# 108415), APC anti-mouse IgM (Biolegend, Cat# 406509), APC-eFluor 780 anti-mouse Mac1 (eBioscience, Cat# 47-0112-82), PerCP/Cy5.5 anti-mouse Sca-1 (Biolegend, Cat# 108123), BV510 anti-mouse Ter119 (Biolegend, Cat# 116237), FITC anti-mouse CD2 (eBioscience, 11-0021-81), FITC anti-mouse CD3 (Biolegend, Cat# 100204), FITC anti-mouse CD5 (Biolegend, Cat# 100606), FITC anti-mouse CD8a (Biolegend, Cat# 100706), FITC anti-mouse B220 (eBioscience, 11-0452-85), FITC anti-mouse Gr-1 (Biolegend, Cat# 108406), FITC anti-mouse Ter119 (Biolegend, Cat# 116206), PE/Cy7 Streptavidin (Biolegend, Cat# 405206).

### RNA-seq and Data Analysis

RNA-seq library was prepared using the Ovation RNA-seq system (NuGEN). Sequencing reads from all RNA-seq experiments were aligned to human (hg19) reference genome by STAR 2.5.2b ^66^ with the parameters: --outFilterMultimapNmax 1. RSEM was used to calculate normalized gene expression (Transcripts per Million Reads or TPM) ^67^. Differential gene expression analysis was performed by DESeq2 ^68^.

### ChIP-seq and Data Analysis

ChIP-seq was performed as described ^35^ using antibodies for HA (Santa Cruz Biotechnology, sc-805), Cas9 (Abcam, ab191468), H3K4me1 (Abcam, ab8895), H3K4me2 (Millipore, 07-030), H3K27ac (Abcam, ab4729), H3K9me3 (Abcam, ab8898), GATA1 (Abcam, ab11852), TAL1 (Santa Cruz Biotechnology, sc-12984), and CTCF (Millipore, 07-729) in HEK293T and/or K562 cells, respectively. Briefly, cross-linked chromatin was sonicated in RIPA 0 buffer (10mM Tris-HCl, 1mM EDTA, 0.1% sodium deoxycholate, 0.1% SDS, 1% Triton X-100, 0.25% Sarkosyl, pH8.0) to 200∼500bp. Final concentration 150mM NaCl was added to the chromatin and antibody mixture before incubation overnight at 4°C. Chromatin was washed and ChIP DNA was purified. ChIP-seq libraries were generated using NEBNext Ultra II DNA library prep kit following the manufacturer’s protocol (NEB), and sequenced on an Illumina NextSeq500 system using the 75bp high output sequencing kit. ChIP-seq raw reads were aligned to the human hg19 genome assembly using Bowtie2 ^69^ with the default parameters. Only tags that uniquely mapped to the genome were used for further analysis. ChIP-seq peaks were called by MACS using the “--nomodel” parameter ^70^. Peaks that overlap with the ENCODE blacklist regions ^3^ or the validated non-targeting sgRNA (sgGal4) enriched regions (chr6:74,229,700-74,231,800; chr3:17,443,100-17,444,200 and chr15:68,131,900-68,133,000) were removed. To compare ChIP-seq signal intensities in samples prepared from cells expressing the target-specific sgRNAs versus the non-targeting sgGal4, MAnorm ^71^ was applied to remove systematic bias between samples and then calculate the normalized ChIP-seq read densities of each peak for all samples. The window size was 1000bp which matched the average width of the identified ChIP-seq peaks.

### *In Vivo* Enhancer Perturbation Screen and Data Analysis

CD45.2^+^ BM lineage negative HSPCs from dCas9-KRAB KI mice were isolated by LS columns (Miltenyi Biotec) and cultured in S-clone SF-O3 medium (Iwai North America) containing 1% fetal bovine serum (FBS), 1% penicillin/streptomycin (P/S), 1% supplement, 50µM β-Mercaptoethanol, 50ng/ml mouse SCF and 50ng/ml mouse TPO. After overnight culture, cells were transduced with retroviruses containing target-specific or non-targeting control sgRNAs at MOI ≤ 0.5. Twenty million cells per mouse were transplanted into CD45.1^+^ lethally irradiated C57BL/6 recipient mice, followed by Dox (2mg/ml, supplemented with sucrose at 10mg/ml) administration in drinking water to induce dCas9-KRAB expression for 16 weeks. By this time, all donor-derived hematopoietic cells (CD45.2^+^) were differentiated from transplanted donor HSCs instead of short-lived progenitors. We collected cells before bone marrow transplantation (T1) as the baseline control and the GFP positive donor-derived mature myeloid (Gr1^+^Mac1^+^), B (B220^+^) and T (CD3^+^) lymphoid cells in the peripheral blood of recipients 16 weeks post-transplantation (T2). Genomic DNA was isolated and sgRNA sequences were PCR amplified using primers listed in Table S2. PCR amplicon libraries were generated using NEBNext Ultra II DNA library prep kit following the manufacturer’s protocol (NEB), and sequenced on an Illumina NextSeq500 system using the 75bp high output sequencing kit. Two biological replicate experiments for each enCRISPRi screen were performed. In each replicate experiment, myeloid, B and T cells were isolated from 3 (for locus-specific perturbation) or 15 (for multi-loci perturbation) independent recipient mice, and genomic DNA were isolated and pooled for the sgRNA amplicon sequencing analysis. After sequencing, the sgRNA sequences were extracted from raw fastq files and mapped to sgRNA sequences in enCRISPRi screen libraries. Reads of each sgRNA were counted and normalized to the total read counts of each sample. Mean sgRNA counts from replicates were calculated at starting (T1) and end time point (T2). The growth phenotype of each sgRNA was quantified as log2 transformed mean sgRNA count ratio between T2 and T1. For calculation of the significance (*P* value) of depletion for each targeted enhancer, promoter or non-targeting control (NC) regions in each cell type, we used MAGECK ^72^ test with the parameters of --norm-method total --gene-lfc-method mean.

## QUANTIFICATION AND STATISTICAL ANALYSIS

Statistical details including *N*, mean and statistical significance values are indicated in the text, figure legends, or methods. Error bars in the experiments represent standard error of the mean (SEM) or standard deviation (SD) from either independent experiments or independent samples. All statistical analyses were performed using GraphPad Prism, and the detailed information about statistical methods is specified in figure legends or methods.

## REPORTING SUMMARY

Further information on experimental design is available in the Nature Research Reporting Summary linked to this article.

## DATA AND SOFTWARE AVAILABILITY

All raw and processed ATAC-seq, ChIP-seq and RNA-seq data are available in the Gene Expression Omnibus (GEO): GSE132216. Computation codes are available from the corresponding author on request.

