## Supplemental Information for "Interrogation of Enhancer Function by Enhanced CRISPR Epigenetic Editing"

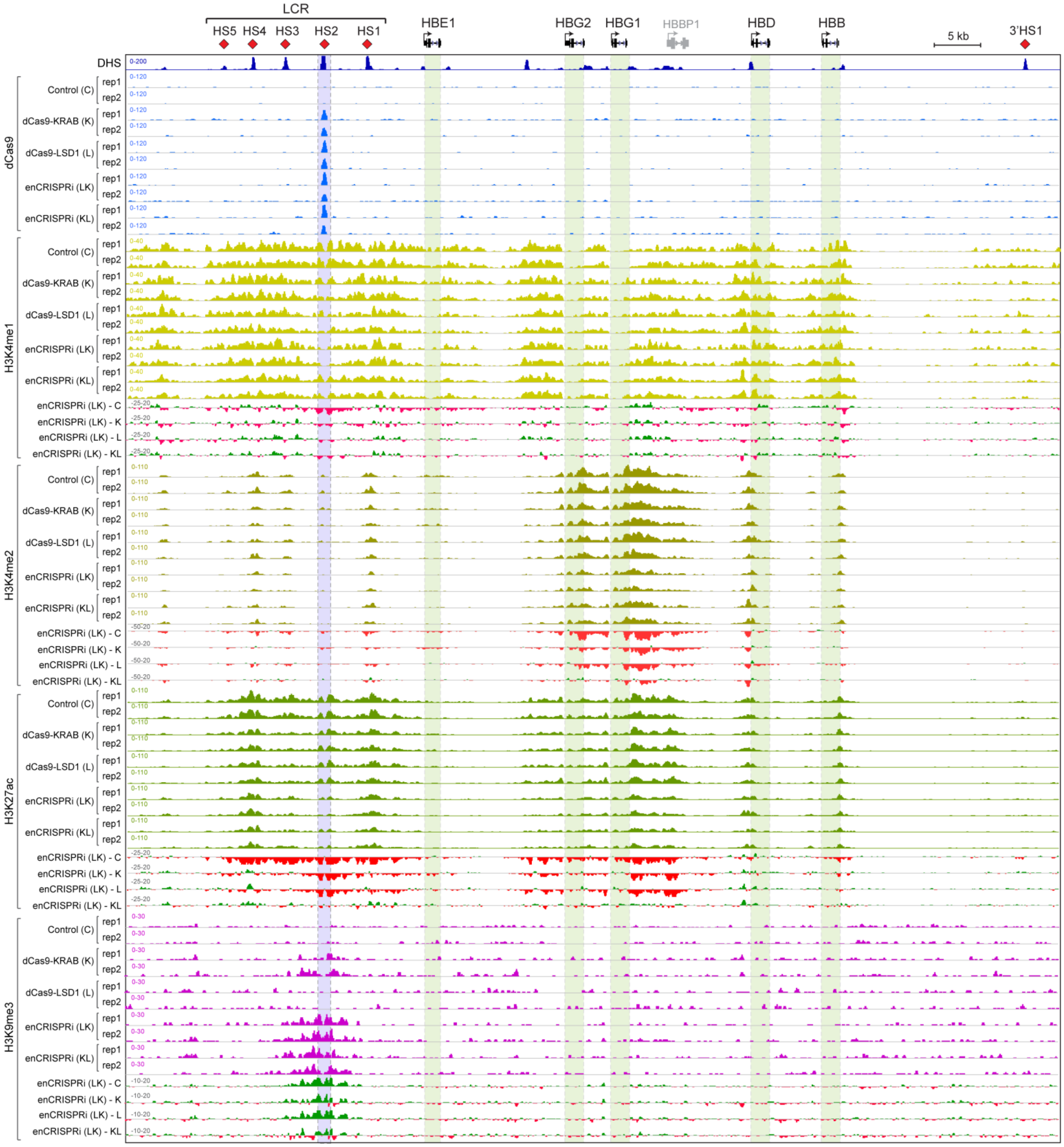


**Figure S1. Targeting enCRISPRi to the β-Globin Gene Cluster Leads to Epigenetic Reprogramming**

Similar to Fig. 3, density maps are shown for ChIP-seq of dCas9, H3K4me1, H3K4me2, H3K27ac, and H3K9me3 at the β-globin cluster (chr11: 5,222,500-5,323,700; hg19) in K562 cells co-expressing non-targeting sgGal4 (control or C) or sgHS2 with dCas9-KRAB (K), dCas9-LSD1 (L) or enCRISPRi (LK and KL). Regions showing increased or decreased ChIP-seq signals in enCRISPRi (LK) relative to control, dCas9-KRAB, dCas9-LSD1 or enCRISPRi (KL) (enCRISPRi – C, enCRISPRi – K, enCRISPRi – L, or enCRISPRi – KL) are depicted in *green* and *red*, respectively. Blue bars denote the sgRNA-targeted HS2 enhancer. Green bars denote the β-globin genes. Independent replicates (rep1 and rep2) are shown for all ChIP-seq experiments. For side-by-side comparisons, the same scales were used for ChIP-seq datasets by the same antibody in different conditions.

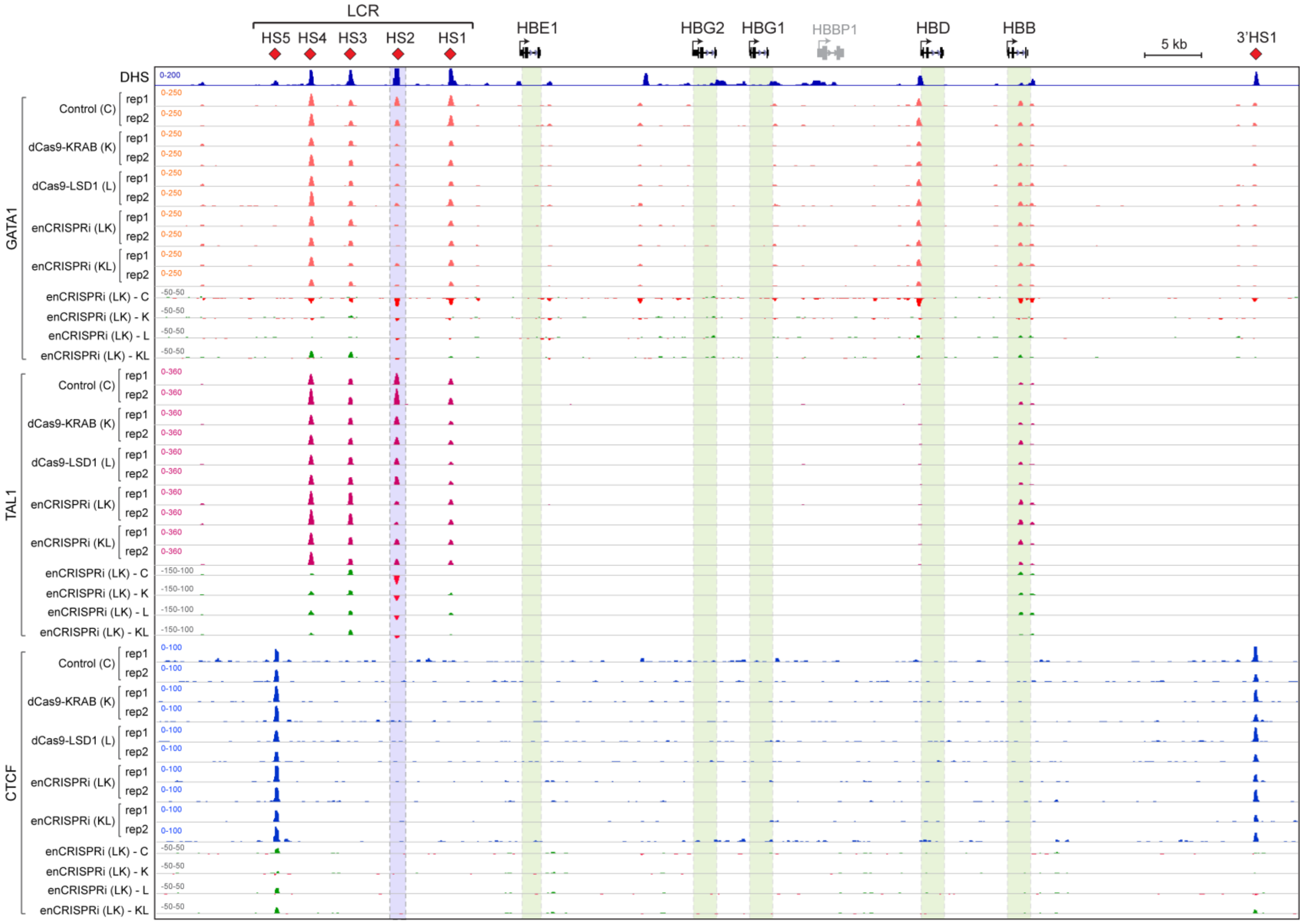


**Figure S2. Targeting enCRISPRi to the β-Globin Gene Cluster Interferes with TF Binding**
Similar to Fig. 3, density maps are shown for ChIP-seq of GATA1, TAL1, and CTCF at the β-globin cluster (chr11: 5,222,500-5,323,700; hg19) in K562 cells. Independent ChIP-seq replicates (rep1 and rep2) are shown. For side-by-side comparisons, the same scales were used for ChIP-seq datasets by the same antibody in different conditions.

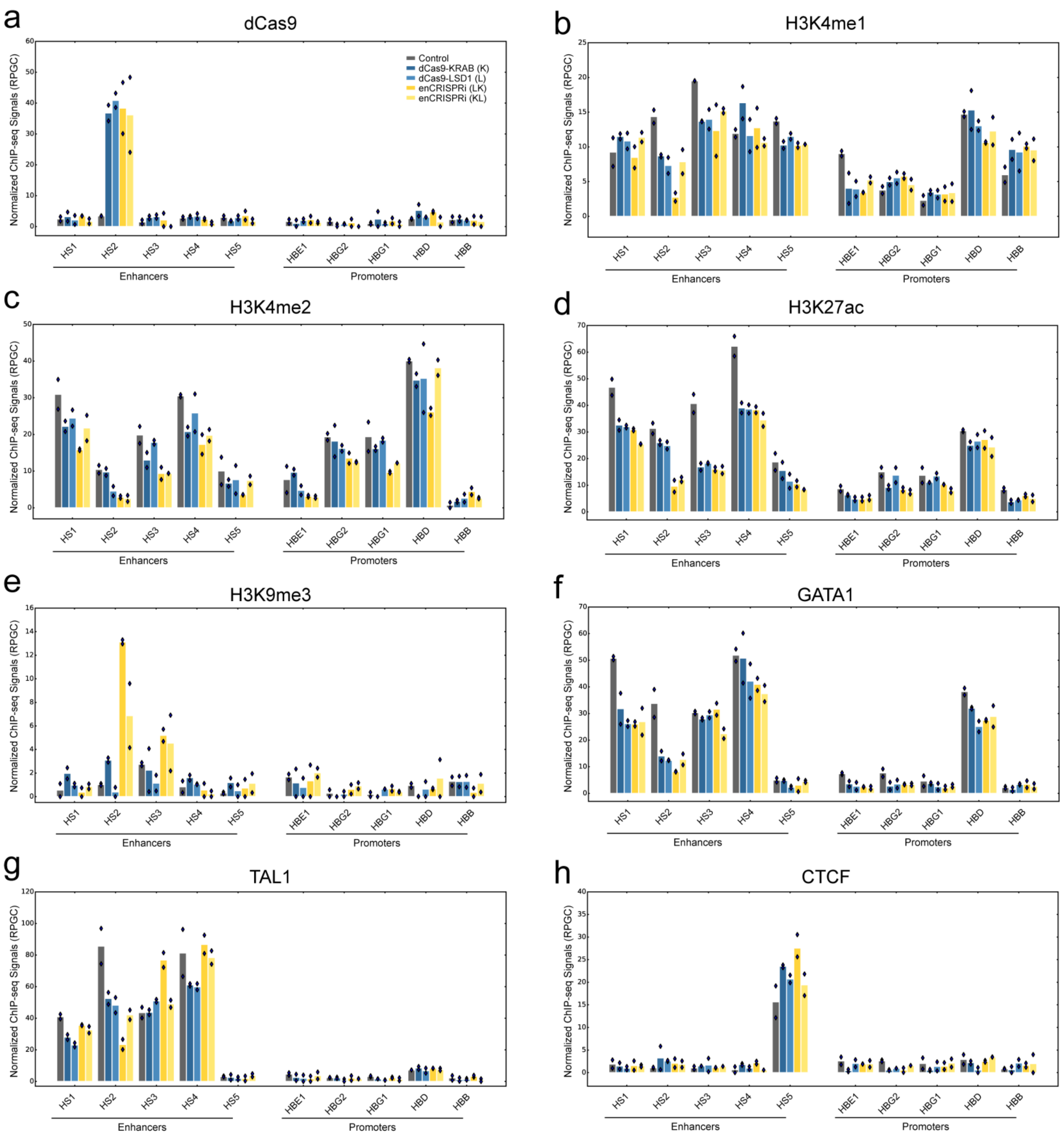


**Figure S3. Analysis of ChIP-seq Signals at the β-Globin Enhancers and Promoters**

Bargraphs are shown for the normalized ChIP-seq signals as reads per genomic content (PRGC) at the β-globin enhancers (HS1 to HS5) and promoters (HBE1, HBG2, HBG1, HBD and HBB) for dCas9 (a), H3K4me1 (b), H3K4me2 (c), H3K27ac (d), H3K9me3 (e), GATA1 (f), TAL1 (g), and CTCF (h) in K562 cells co-expressing non-targeting sgGal4 (control) or sgHS2 with dCas9-KRAB (K), dCas9-LSD1 (L) or enCRISPRi (LK and KL). Dots indicate values from the independent ChIP-seq replicates (rep1 and rep2).

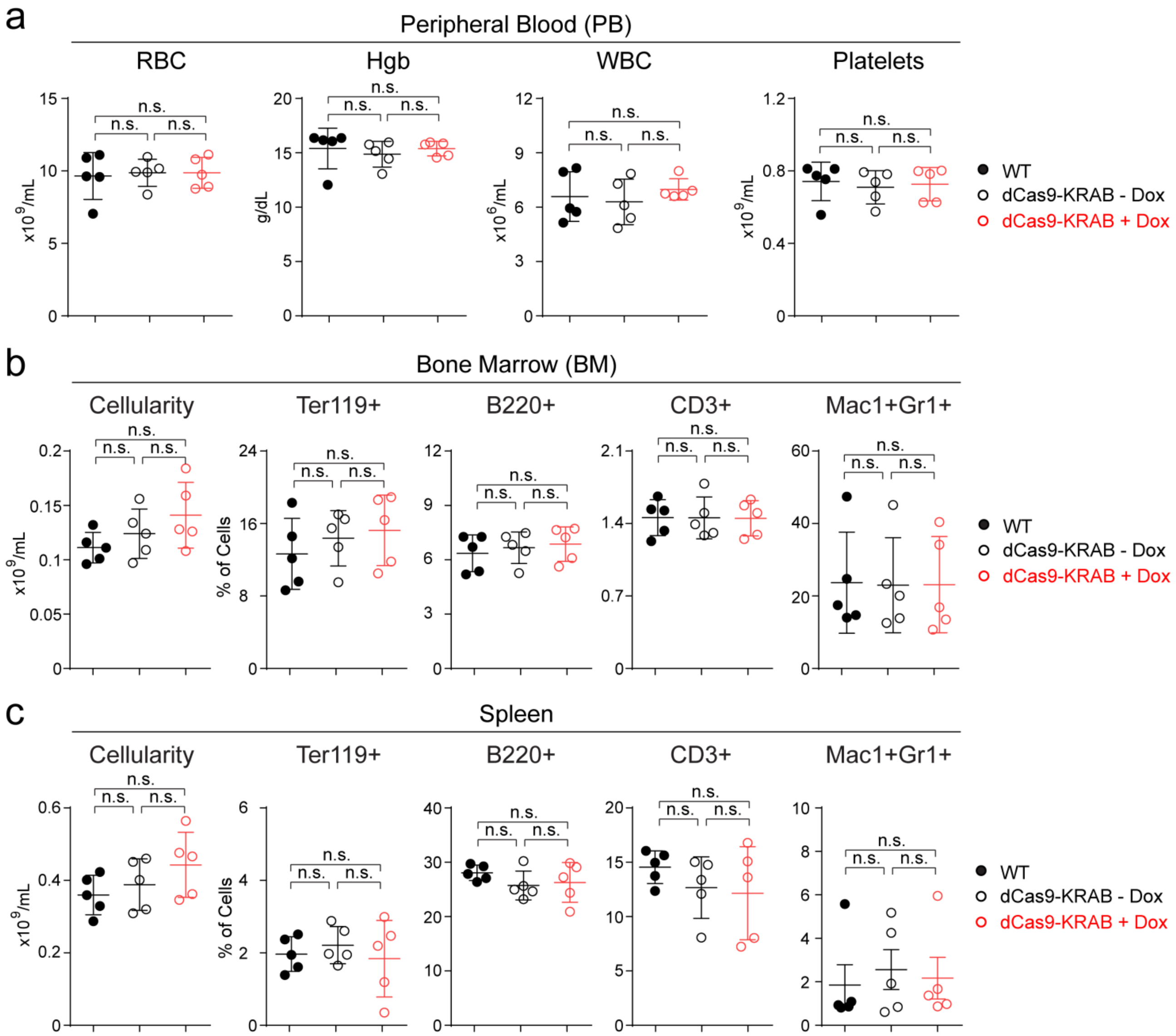


**Figure S4. Inducible dCas9-KRAB Expression Had No Effect on Mature Hematopoietic Cell Types**

**(a)** Complete blood count (CBC) of peripheral blood (PB) red blood cells (RBC), hemoglobin (Hgb), white blood cells (WBC) and platelets in mice 8 weeks after Dox treatment. *N* = 5 mice per group. Results are mean ± SD and analyzed by a repeated-measures one-way ANOVA with multiple comparisons. n.s. not significant.

**(b)** BM cellularity and frequencies of erythroid ($Ter119^+$), B-lymphoid ($B220^+$), T-lymphoid ($CD3^+$) and myeloid ($Mac1^+Gr1^+$) cells in mice 8 weeks after Dox treatment.

**(c)** Spleen cellularity and frequencies of erythroid, B-lymphoid, T-lymphoid and myeloid cells 8 weeks after Dox treatment.

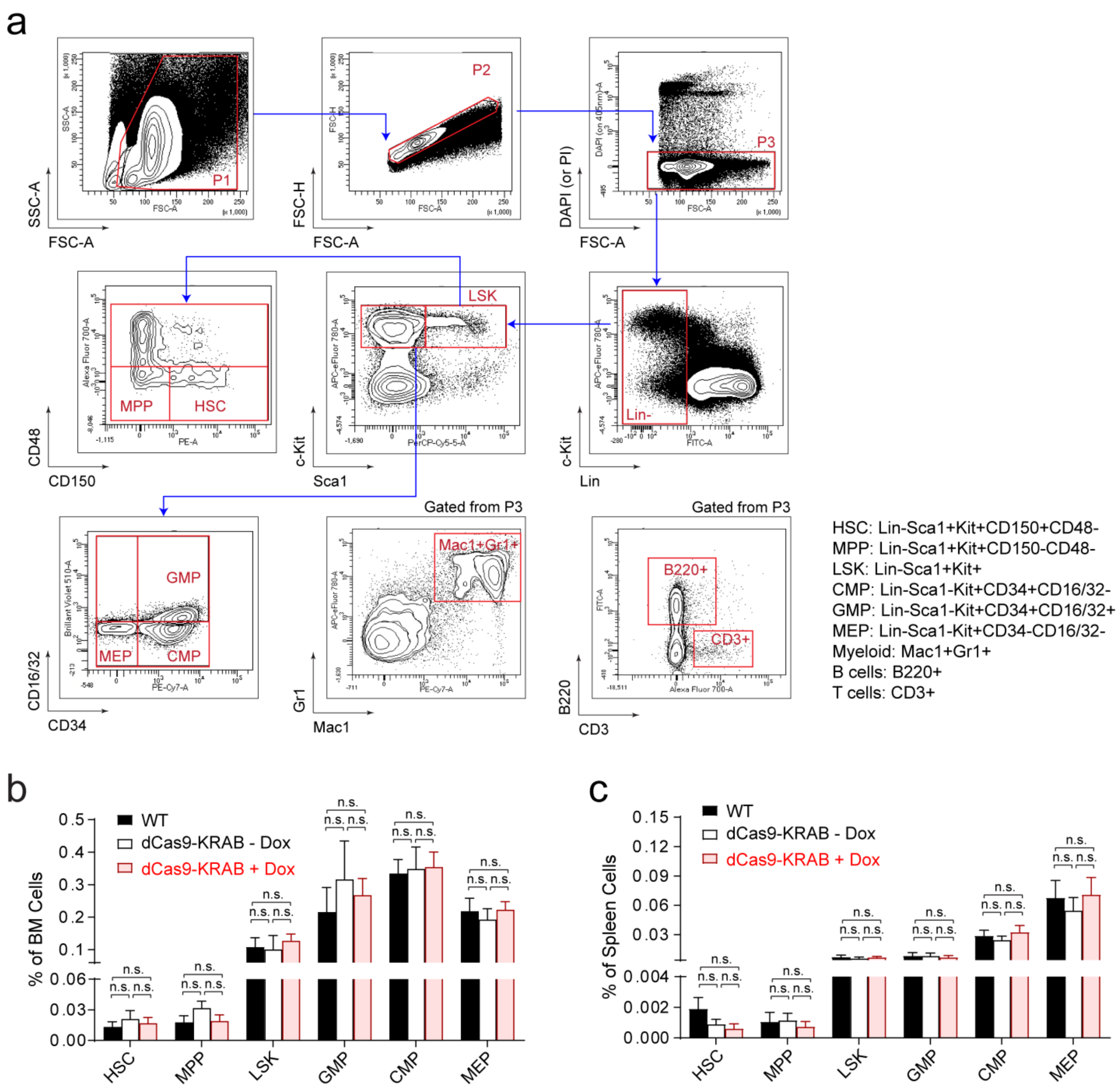


**Figure S5. Inducible dCas9-KRAB Expression Had No Effect on Hematopoietic Differentiation**

**(a)** Representative flow cytometry gates are shown for the analysis of various hematopoietic stem/progenitor cells (HSPCs) and mature lineages in mouse bone marrow.

**(b)** Frequencies of HSPCs in BM 8 weeks after Dox treatment. *N* = 5 mice per group. Results are mean ± SD and analyzed by a one-way ANOVA with multiple comparisons. n.s. not significant.

**(c)** Frequencies of HSPCs in spleen 8 weeks after Dox treatment. *N* = 5 mice per group.

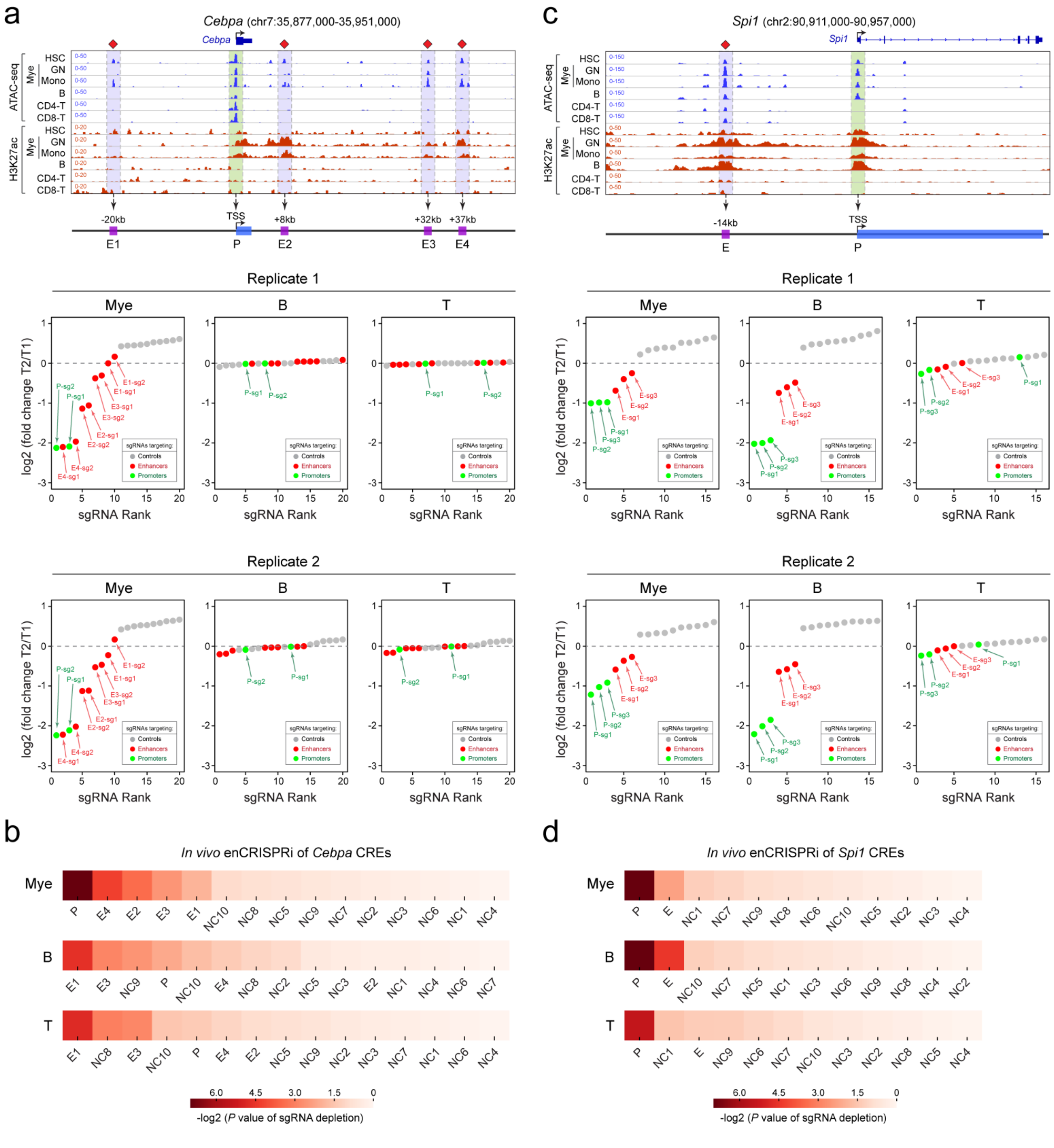

**Figure S6. Locus-Specific Enhancer Perturbation at *Cebpa* and *Spi1* Loci**

**(a)** *In vivo* enCRISPRi perturbation of *Cebpa* CREs during hematopoiesis. Similar to Fig. 5e, waterfall plots are shown for target-specific sgRNAs (*green* and *red* dots) and non-targeting control sgRNAs (*grey* dots) by the mean normalized log2 fold changes in myeloid, T or B cells 16-weeks post-BMT (T2) relative to pooled sgRNA-transduced HSPCs (T1) in two independent replicate screens ($N$ = 3 recipient mice per screen). The annotated *Cebpa* promoter (P) and enhancers (E1 to E4) are indicated by *green* and *blue* shaded lines.

**(b)** Ranking of the top depleted *Cebpa* CREs by the –log2 (*P* value of sgRNA depletion). The significance (*P* value) of depletion for each targeted enhancer, promoter or non-targeting control (NC) region in each cell type was calculated using the MAGECK test. P, promoter; E, enhancer; NC, non-targeting control.

**(c)** *In vivo* enCRISPRi perturbation of *Spi1* CREs during hematopoiesis. Similar to Fig. 5f, waterfall plots are shown for two independent replicate screens (*N* = 3 recipient mice per screen). The annotated *Spi1* promoter (P) and enhancer (E) are indicated by *green* and *blue* shaded lines.

**(d)** Ranking of the top depleted *Spi1* CREs by the –log2 (*P* value of sgRNA depletion).

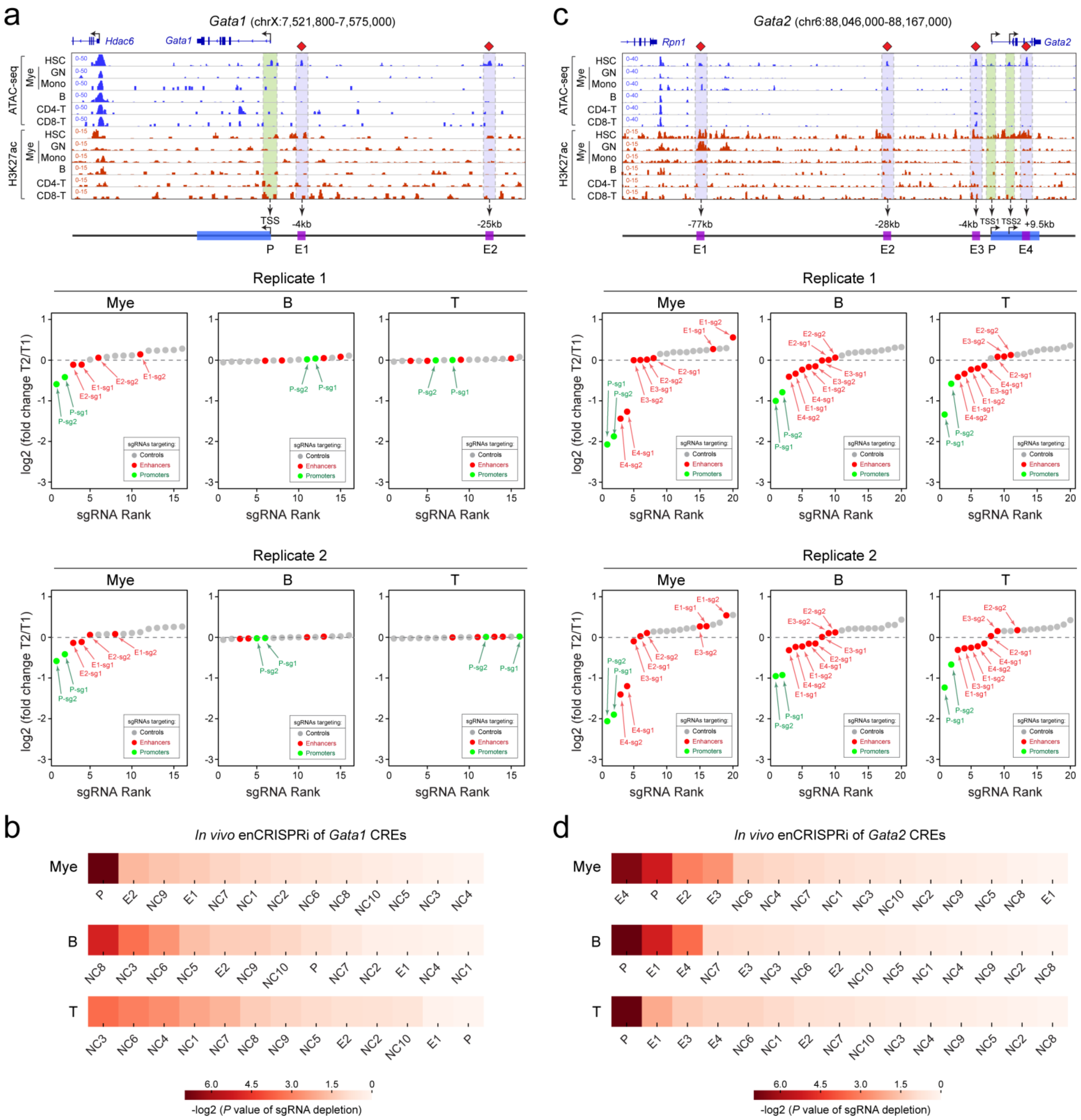


**Figure S7. Locus-Specific Enhancer Perturbation at *Gata1* and *Gata2* Loci**

**(a)** *In vivo* enCRISPRi perturbation of *Gata1* CREs during hematopoiesis. Density maps are shown for ATAC-seq and H3K27ac ChIP-seq at the *Gata1* locus (chrX:7,521,800-7,575,000; mm9) in bone marrow HSC, granulocytes (GN), monocytes (Mono), B, CD4+ and CD8+ T cells, respectively. Waterfall plots are shown for target-specific sgRNAs (*green* and *red* dots) and non-targeting control sgRNAs (*grey* dots) by the mean normalized log2 fold changes in myeloid, T or B cells 16-weeks post-BMT (T2) relative to pooled sgRNA-transduced HSPCs (T1) in two independent replicate screens (*N* = 3 recipient mice per screen). The annotated *Gata1* promoter (P) and enhancers (E1 and E2) are indicated by *green* and *blue* shaded lines.

**(b)** Ranking of the top depleted *Gata1* CREs by the –log2 (*P* value of sgRNA depletion). The significance (*P* value) of depletion for each targeted enhancer, promoter or non-targeting control (NC) region in each cell type was calculated using the MAGECK test. P, promoter; E, enhancer; NC, non-targeting control.

**(c)** *In vivo* enCRISPRi perturbation of *Gata2* CREs during hematopoiesis. Density maps are shown for ATAC-seq and H3K27ac ChIP-seq at the *Gata2* locus (chr6:88,046,000-88,167,000; mm9) in bone marrow HSC, GN, Mono, B, CD4+ and CD8+ T cells, respectively. Waterfall plots are shown for two independent replicate screens (*N* = 3 recipient mice per screen). The annotated *Gata2* promoter (P) and enhancers (E1 to E4) are indicated by *green* and *blue* shaded lines.

**(d)** Ranking of the top depleted *Gata2* CREs by the –log2 (*P* value of sgRNA depletion).

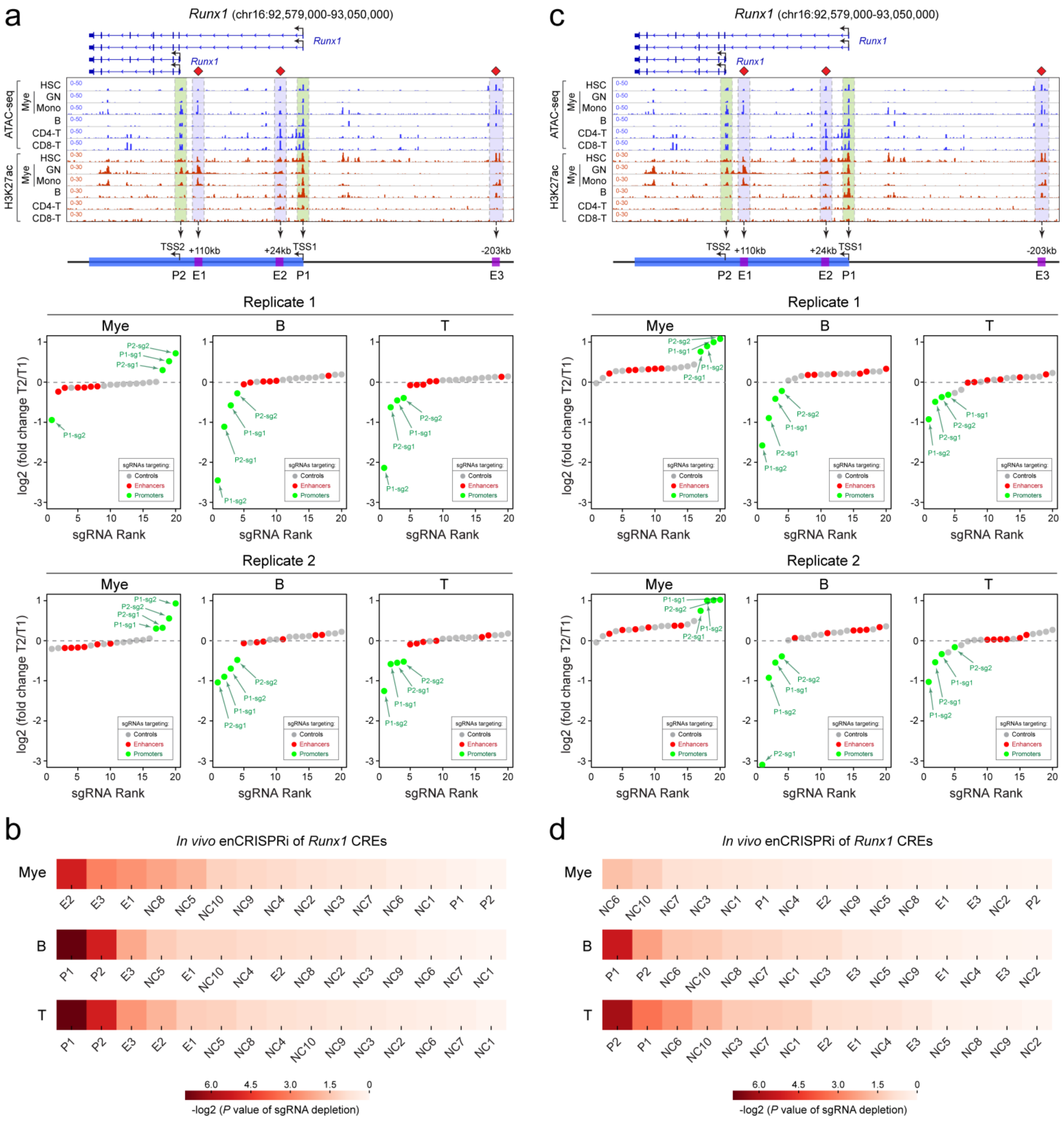


### Figure S8. Locus-Specific and Multi-loci Perturbations of *Runx1* CREs

**(a)** *In vivo* enCRISPRi perturbation of *Runx1* CREs by locus-specific enCRISPRi screen during hematopoiesis. Similar to Fig. 6c, waterfall plots are shown for two independent replicate screens (*N* = 3 recipient mice per screen). The annotated *Runx1* promoters (P1 and P2) and enhancers (E1 to E3) are indicated by *green* and *blue* shaded lines.

**(b)** Ranking of the top depleted *Runx1* CREs by the –log2 (*P* value of sgRNA depletion) by locus-specific enCRISPRi screen. The significance (*P* value) of depletion for each targeted enhancer, promoter or non-targeting control (NC) region in each cell type was calculated using the MAGECK test. P, promoter; E, enhancer; NC, non-targeting control.

**(c)** *In vivo* enCRISPRi perturbation of *Runx1* CREs by multi-loci enCRISPRi screen during hematopoiesis. Similar to Fig. 6d, waterfall plots are shown for two independent replicate screens (*N* = 3 recipient mice per experiment).

**(d)** Ranking of the top depleted *Runx1* CREs by the –log2 (*P* value of sgRNA depletion) in the multi-loci enCRISPRi screen.

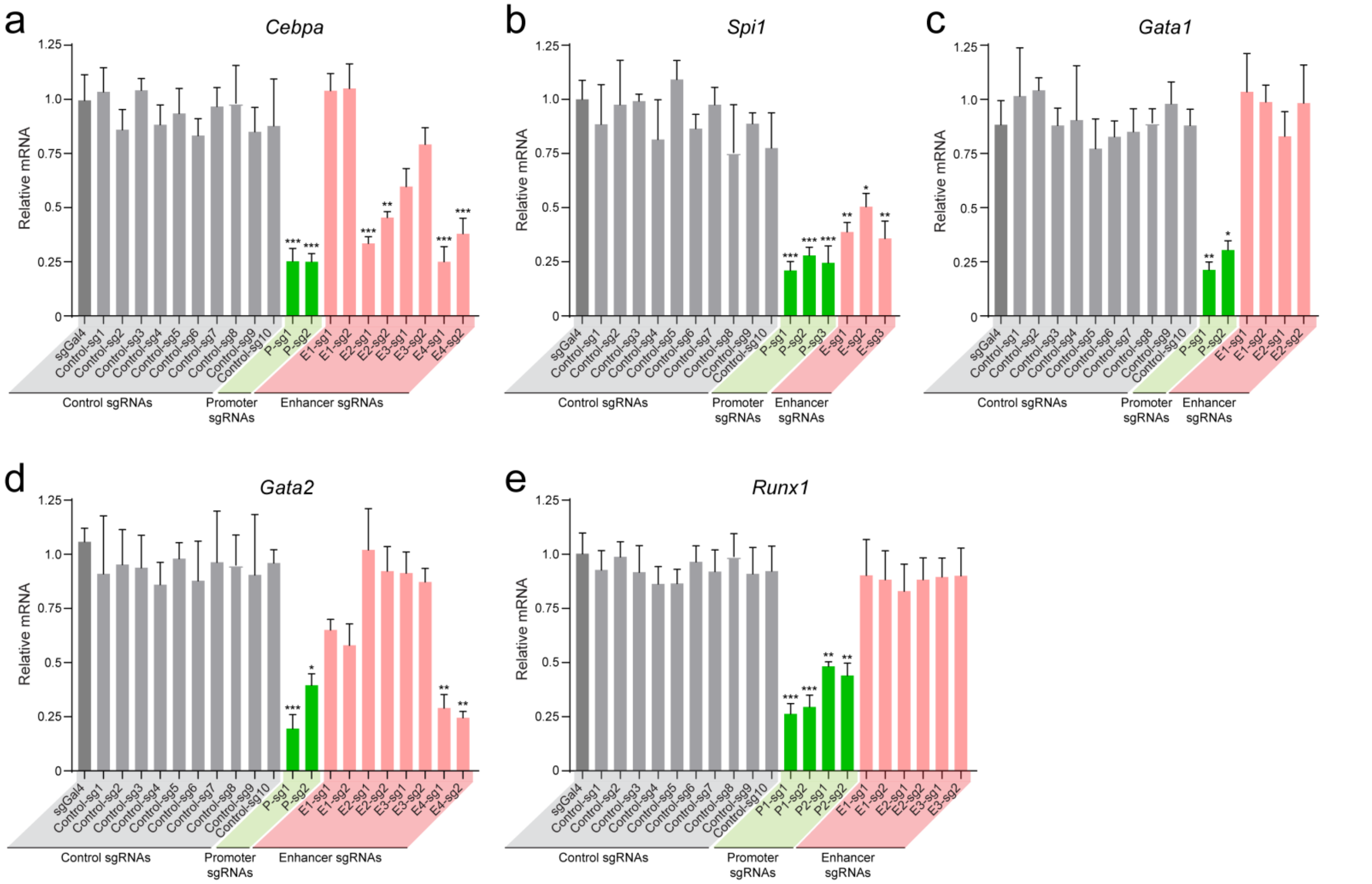


**Figure S9. *In Vivo* enCRISPRi of Promoters and Enhancers Impairs Gene Expression**

**(a)** enCRISPRi-mediated perturbation of *Cebpa* CREs impaired its mRNA expression in BM HSPCs. Independent sgRNAs (sg1 and sg2) for each targeted enhancer or promoter are shown. Cells transduced with non-targeting sgGal4 were analyzed as the control. Results are mean ± SEM and analyzed by a one-way ANOVA. *$P < 0.05$, **$P < 0.01$, ***$P < 0.001$, n.s. not significant.

**(b)** Expression of *Spi1* mRNA upon enCRISPRi-mediated perturbation of CREs in HSPCs.

**(c)** Expression of *Gata1* mRNA upon enCRISPRi-mediated perturbation of CREs in HSPCs.

**(d)** Expression of *Gata2* mRNA upon enCRISPRi-mediated perturbation of CREs in HSPCs.

**(e)** Expression of *Runx1* mRNA upon enCRISPRi-mediated perturbation of CREs in HSPCs.

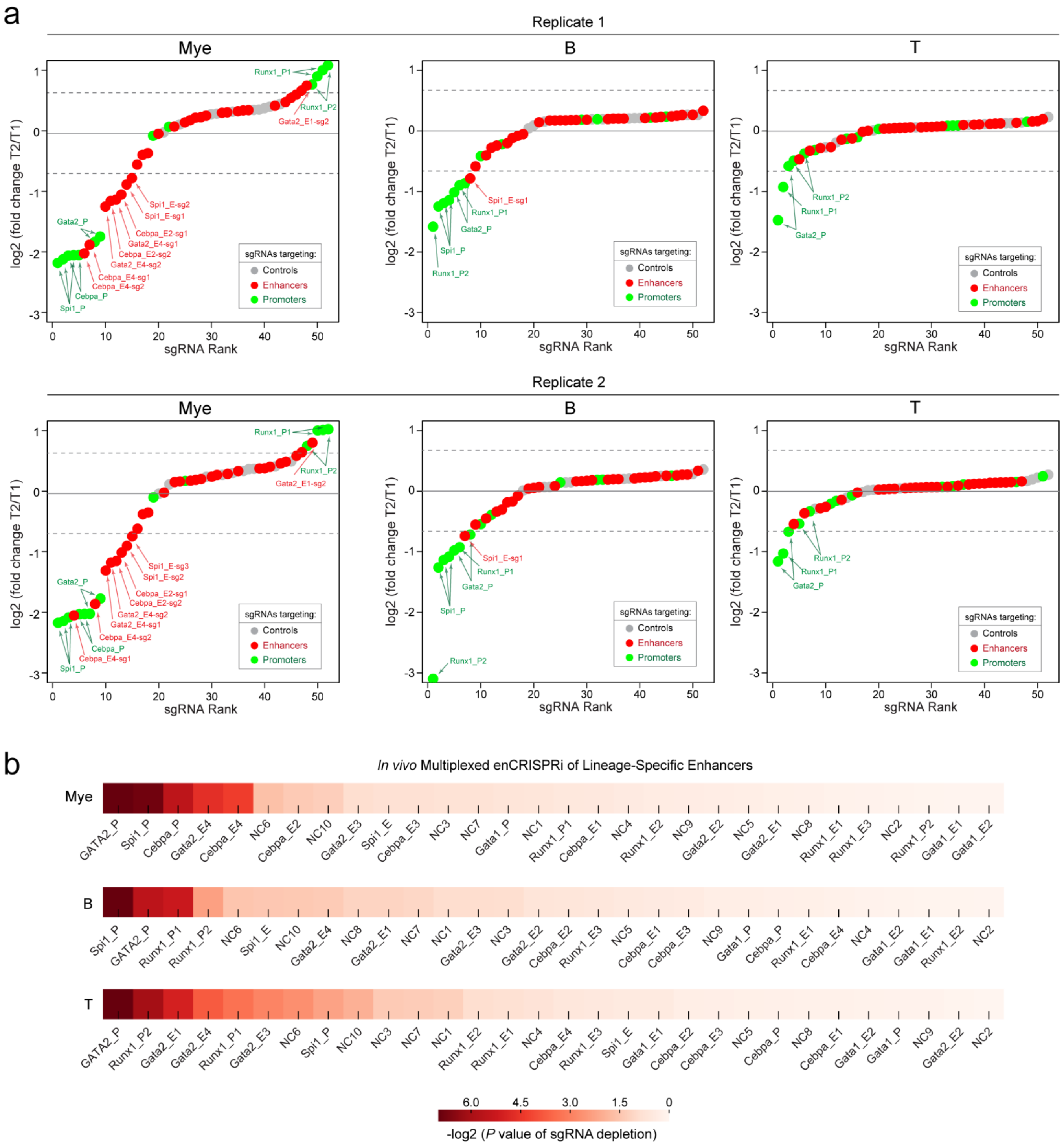


**Figure S10. Multiplexed *In Vivo* enCRISPRi Perturbations of CREs during Hematopoiesis**

**(a)** *In vivo* perturbation of annotated CREs for five key hematopoietic TFs. Similar to Fig. 6b, waterfall plots are shown for two independent replicate screens ($N$ = 15 recipient mice per screen).

**(b)** Ranking of the top depleted CREs by the –log2 ($P$ value of sgRNA depletion) by multiplexed enCRISPRi screens. The significance ($P$ value) of depletion for each targeted enhancer, promoter or non-targeting control (NC) region in each cell type was calculated using the MAGECK test. P, promoter; E, enhancer; NC, non-targeting control.

### SUPPLEMENTARY TABLES

#### Table S1. List of Genomic Datasets Used in This Study

The name, data type, cell type, GEO accession number and citation for each dataset are shown.

#### Table S2. Sequences of Primers and sgRNAs

The name and sequence of each primer or sgRNA are shown.

***Table S1. List of Genomic Datasets Used in This Study***

| ***Datasets*** | ***Data Type*** | ***Cell Type*** | ***GEO ID*** | ***Citation*** |
|---|---|---|---|---|
| ChIP-seq_Jurkat-H3K27ac | ChIP-seq | Jurkat | GSM3854054 | This study |
| ChIP-seq_293T_enCRISPRa_sgGal4_HA_rep1 | ChIP-seq | 293T | GSM3854055 | This study |
| ChIP-seq_293T_enCRISPRa_sgGal4_HA_rep2 | ChIP-seq | 293T | GSM3854056 | This study |
| ChIP-seq_293T_enCRISPRa_sgHS2_HA_rep1 | ChIP-seq | 293T | GSM3854057 | This study |
| ChIP-seq_293T_enCRISPRa_sgHS2_HA_rep2 | ChIP-seq | 293T | GSM3854058 | This study |
| ChIP-seq_K562_enCRISPRi-LK_sgGal4_HA_rep1 | ChIP-seq | K562 | GSM3854059 | This study |
| ChIP-seq_K562_enCRISPRi-LK_sgGal4_HA_rep2 | ChIP-seq | K562 | GSM3854060 | This study |
| ChIP-seq_K562_enCRISPRi-LK_sgHS2_HA_rep1 | ChIP-seq | K562 | GSM3854061 | This study |
| ChIP-seq_K562_enCRISPRi-LK_sgHS2_HA_rep2 | ChIP-seq | K562 | GSM3854062 | This study |
| ChIP-seq_K562_dCas9-KRAB_sgHS2_cas9_rep1 | ChIP-seq | K562 | GSM3854063 | This study |
| ChIP-seq_K562_dCas9-KRAB_sgHS2_cas9_rep2 | ChIP-seq | K562 | GSM3854064 | This study |
| ChIP-seq_K562_dCas9-KRAB_sgHS2_CTCF_rep1 | ChIP-seq | K562 | GSM3854065 | This study |
| ChIP-seq_K562_dCas9-KRAB_sgHS2_CTCF_rep2 | ChIP-seq | K562 | GSM3854066 | This study |
| ChIP-seq_K562_dCas9-KRAB_sgHS2_GATA1_rep1 | ChIP-seq | K562 | GSM3854067 | This study |
| ChIP-seq_K562_dCas9-KRAB_sgHS2_GATA1_rep2 | ChIP-seq | K562 | GSM3854068 | This study |
| ChIP-seq_K562_dCas9-KRAB_sgHS2_H3K27ac_rep1 | ChIP-seq | K562 | GSM3854069 | This study |
| ChIP-seq_K562_dCas9-KRAB_sgHS2_H3K27ac_rep2 | ChIP-seq | K562 | GSM3854070 | This study |
| ChIP-seq_K562_dCas9-KRAB_sgHS2_H3K4me2_rep1 | ChIP-seq | K562 | GSM3854071 | This study |
| ChIP-seq_K562_dCas9-KRAB_sgHS2_H3K4me2_rep2 | ChIP-seq | K562 | GSM3854072 | This study |
| ChIP-seq_K562_dCas9-KRAB_sgHS2_H3K4me_rep1 | ChIP-seq | K562 | GSM3854073 | This study |
| ChIP-seq_K562_dCas9-KRAB_sgHS2_H3K4me_rep2 | ChIP-seq | K562 | GSM3854074 | This study |
| ChIP-seq_K562_dCas9-KRAB_sgHS2_H3K9me3_rep1 | ChIP-seq | K562 | GSM3854075 | This study |
| ChIP-seq_K562_dCas9-KRAB_sgHS2_H3K9me3_rep2 | ChIP-seq | K562 | GSM3854076 | This study |
| ChIP-seq_K562_dCas9-KRAB_sgHS2_TAL1_rep1 | ChIP-seq | K562 | GSM3854077 | This study |
| ChIP-seq_K562_dCas9-KRAB_sgHS2_TAL1_rep2 | ChIP-seq | K562 | GSM3854078 | This study |
| ChIP-seq_K562_dCas9-LSD1_sgHS2_cas9_rep1 | ChIP-seq | K562 | GSM3854079 | This study |
| ChIP-seq_K562_dCas9-LSD1_sgHS2_cas9_rep2 | ChIP-seq | K562 | GSM3854080 | This study |
| ChIP-seq_K562_dCas9-LSD1_sgHS2_CTCF_rep1 | ChIP-seq | K562 | GSM3854081 | This study |
| ChIP-seq_K562_dCas9-LSD1_sgHS2_CTCF_rep2 | ChIP-seq | K562 | GSM3854082 | This study |
| ChIP-seq_K562_dCas9-LSD1_sgHS2_GATA1_rep1 | ChIP-seq | K562 | GSM3854083 | This study |
| ChIP-seq_K562_dCas9-LSD1_sgHS2_GATA1_rep2 | ChIP-seq | K562 | GSM3854084 | This study |
| ChIP-seq_K562_dCas9-LSD1_sgHS2_H3K27ac_rep1 | ChIP-seq | K562 | GSM3854085 | This study |
| ChIP-seq_K562_dCas9-LSD1_sgHS2_H3K27ac_rep2 | ChIP-seq | K562 | GSM3854086 | This study |
| ChIP-seq_K562_dCas9-LSD1_sgHS2_H3K4me2_rep1 | ChIP-seq | K562 | GSM3854087 | This study |
| ChIP-seq_K562_dCas9-LSD1_sgHS2_H3K4me2_rep2 | ChIP-seq | K562 | GSM3854088 | This study |
| ChIP-seq_K562_dCas9-LSD1_sgHS2_H3K4me_rep1 | ChIP-seq | K562 | GSM3854089 | This study |
| ChIP-seq_K562_dCas9-LSD1_sgHS2_H3K4me_rep2 | ChIP-seq | K562 | GSM3854090 | This study |
| ChIP-seq_K562_dCas9-LSD1_sgHS2_H3K9me3_rep1 | ChIP-seq | K562 | GSM3854091 | This study |
| ChIP-seq_K562_dCas9-LSD1_sgHS2_H3K9me3_rep2 | ChIP-seq | K562 | GSM3854092 | This study |
| ChIP-seq_K562_dCas9-LSD1_sgHS2_TAL1_rep1 | ChIP-seq | K562 | GSM3854093 | This study |
| ChIP-seq_K562_dCas9-LSD1_sgHS2_TAL1_rep2 | ChIP-seq | K562 | GSM3854094 | This study |
| ChIP-seq_K562_enCRISPRi-KL_sgHS2_cas9_rep1 | ChIP-seq | K562 | GSM3854095 | This study |
| ChIP-seq_K562_enCRISPRi-KL_sgHS2_cas9_rep2 | ChIP-seq | K562 | GSM3854096 | This study |
| ChIP-seq_K562_enCRISPRi-KL_sgHS2_CTCF_rep1 | ChIP-seq | K562 | GSM3854097 | This study |
| ChIP-seq_K562_enCRISPRi-KL_sgHS2_CTCF_rep2 | ChIP-seq | K562 | GSM3854098 | This study |
| ChIP-seq_K562_enCRISPRi-KL_sgHS2_GATA1_rep1 | ChIP-seq | K562 | GSM3854099 | This study |
| ChIP-seq_K562_enCRISPRi-KL_sgHS2_GATA1_rep2 | ChIP-seq | K562 | GSM3854100 | This study |
| ChIP-seq_K562_enCRISPRi-KL_sgHS2_H3K27ac_rep1 | ChIP-seq | K562 | GSM3854101 | This study |
| ChIP-seq_K562_enCRISPRi-KL_sgHS2_H3K27ac_rep2 | ChIP-seq | K562 | GSM3854102 | This study |
| ChIP-seq_K562_enCRISPRi-KL_sgHS2_H3K4me2_rep1 | ChIP-seq | K562 | GSM3854103 | This study |
| ChIP-seq_K562_enCRISPRi-KL_sgHS2_H3K4me2_rep2 | ChIP-seq | K562 | GSM3854104 | This study |
| ChIP-seq_K562_enCRISPRi-KL_sgHS2_H3K4me_rep1 | ChIP-seq | K562 | GSM3854105 | This study |
| ChIP-seq_K562_enCRISPRi-KL_sgHS2_H3K4me_rep2 | ChIP-seq | K562 | GSM3854106 | This study |
| ChIP-seq_K562_enCRISPRi-KL_sgHS2_H3K9me3_rep1 | ChIP-seq | K562 | GSM3854107 | This study |
| ChIP-seq_K562_enCRISPRi-KL_sgHS2_H3K9me3_rep2 | ChIP-seq | K562 | GSM3854108 | This study |
| ChIP-seq_K562_enCRISPRi-KL_sgHS2_TAL1_rep1 | ChIP-seq | K562 | GSM3854109 | This study |
| ChIP-seq_K562_enCRISPRi-KL_sgHS2_TAL1_rep2 | ChIP-seq | K562 | GSM3854110 | This study |
| ChIP-seq_K562_enCRISPRi-LK_sgHS2_cas9_rep1 | ChIP-seq | K562 | GSM3854111 | This study |
| ChIP-seq_K562_enCRISPRi-LK_sgHS2_cas9_rep2 | ChIP-seq | K562 | GSM3854112 | This study |
| ChIP-seq_K562_enCRISPRi-LK_sgHS2_CTCF_rep1 | ChIP-seq | K562 | GSM3854113 | This study |

| | | | | |
|---|---|---|---|---|
| ChIP-seq_K562_enCRISPRi-LK_sgHS2_CTCF_rep2 | ChIP-seq | K562 | GSM3854114 | This study |
| ChIP-seq_K562_enCRISPRi-LK_sgHS2_GATA1_rep1 | ChIP-seq | K562 | GSM3854115 | This study |
| ChIP-seq_K562_enCRISPRi-LK_sgHS2_GATA1_rep2 | ChIP-seq | K562 | GSM3854116 | This study |
| ChIP-seq_K562_enCRISPRi-LK_sgHS2_H3K27ac_rep1 | ChIP-seq | K562 | GSM3854117 | This study |
| ChIP-seq_K562_enCRISPRi-LK_sgHS2_H3K27ac_rep2 | ChIP-seq | K562 | GSM3854118 | This study |
| ChIP-seq_K562_enCRISPRi-LK_sgHS2_H3K4me2_rep1 | ChIP-seq | K562 | GSM3854119 | This study |
| ChIP-seq_K562_enCRISPRi-LK_sgHS2_H3K4me2_rep2 | ChIP-seq | K562 | GSM3854120 | This study |
| ChIP-seq_K562_enCRISPRi-LK_sgHS2_H3K4me_rep1 | ChIP-seq | K562 | GSM3854121 | This study |
| ChIP-seq_K562_enCRISPRi-LK_sgHS2_H3K4me_rep2 | ChIP-seq | K562 | GSM3854122 | This study |
| ChIP-seq_K562_enCRISPRi-LK_sgHS2_H3K9me3_rep1 | ChIP-seq | K562 | GSM3854123 | This study |
| ChIP-seq_K562_enCRISPRi-LK_sgHS2_H3K9me3_rep2 | ChIP-seq | K562 | GSM3854124 | This study |
| ChIP-seq_K562_enCRISPRi-LK_sgHS2_TAL1_rep1 | ChIP-seq | K562 | GSM3854125 | This study |
| ChIP-seq_K562_enCRISPRi-LK_sgHS2_TAL1_rep2 | ChIP-seq | K562 | GSM3854126 | This study |
| ChIP-seq_K562_sgGal4_Cas9_rep1 | ChIP-seq | K562 | GSM3854127 | This study |
| ChIP-seq_K562_sgGal4_Cas9_rep2 | ChIP-seq | K562 | GSM3854128 | This study |
| ChIP-seq_K562_sgGal4_CTCF_rep1 | ChIP-seq | K562 | GSM3854129 | This study |
| ChIP-seq_K562_sgGal4_CTCF_rep2 | ChIP-seq | K562 | GSM3854130 | This study |
| ChIP-seq_K562_sgGal4_GATA1_rep1 | ChIP-seq | K562 | GSM3854131 | This study |
| ChIP-seq_K562_sgGal4_GATA1_rep2 | ChIP-seq | K562 | GSM3854132 | This study |
| ChIP-seq_K562_sgGal4_H3K27ac_rep1 | ChIP-seq | K562 | GSM3854133 | This study |
| ChIP-seq_K562_sgGal4_H3K27ac_rep2 | ChIP-seq | K562 | GSM3854134 | This study |
| ChIP-seq_K562_sgGal4_H3K4me2_rep1 | ChIP-seq | K562 | GSM3854135 | This study |
| ChIP-seq_K562_sgGal4_H3K4me2_rep2 | ChIP-seq | K562 | GSM3854136 | This study |
| ChIP-seq_K562_sgGal4_H3K4me_rep1 | ChIP-seq | K562 | GSM3854137 | This study |
| ChIP-seq_K562_sgGal4_H3K4me_rep2 | ChIP-seq | K562 | GSM3854138 | This study |
| ChIP-seq_K562_sgGal4_H3K9me3_rep1 | ChIP-seq | K562 | GSM3854139 | This study |
| ChIP-seq_K562_sgGal4_H3K9me3_rep2 | ChIP-seq | K562 | GSM3854140 | This study |
| ChIP-seq_K562_sgGal4_TAL1_rep1 | ChIP-seq | K562 | GSM3854141 | This study |
| ChIP-seq_K562_sgGal4_TAL1_rep2 | ChIP-seq | K562 | GSM3854142 | This study |
| RNA-seq_K562_dCas9-KRAB_sgHS2_rep1 | RNA-seq | K562 | GSM3854143 | This study |
| RNA-seq_K562_dCas9-KRAB_sgHS2_rep2 | RNA-seq | K562 | GSM3854144 | This study |
| RNA-seq_K562_dCas9-LSD1_sgHS2_rep1 | RNA-seq | K562 | GSM3854145 | This study |
| RNA-seq_K562_dCas9-LSD1_sgHS2_rep2 | RNA-seq | K562 | GSM3854146 | This study |
| RNA-seq_K562_enCRISPRi-KL_sgHS2_rep1 | RNA-seq | K562 | GSM3854147 | This study |
| RNA-seq_K562_enCRISPRi-KL_sgHS2_rep2 | RNA-seq | K562 | GSM3854148 | This study |
| RNA-seq_K562_enCRISPRi-LK_sgHS2_rep1 | RNA-seq | K562 | GSM3854149 | This study |
| RNA-seq_K562_enCRISPRi-LK_sgHS2_rep2 | RNA-seq | K562 | GSM3854150 | This study |
| RNA-seq_K562_sgGal4_rep1 | RNA-seq | K562 | GSM3854151 | This study |
| RNA-seq_K562_sgGal4_rep2 | RNA-seq | K562 | GSM3854152 | This study |
| ATAC-seq_Jurkat | ATAC-seq | Jurkat | GSM3854041 | This study |
| ATAC_seq_B | ATAC-seq | B cell | GSM1463172 | Lara-Astiaso et al, Science 2014 |
| ATAC_seq_Monocytes | ATAC-seq | Monocytes | GSM1463174 | Lara-Astiaso et al, Science 2014 |
| ATAC_seq_CD4 | ATAC-seq | T cell | GSM1463175 | Lara-Astiaso et al, Science 2014 |
| ATAC_seq_Granulocytes | ATAC-seq | Granulocytes | GSM1463176 | Lara-Astiaso et al, Science 2014 |
| ATAC_seq_CD8 | ATAC-seq | T cell | GSM1463178 | Lara-Astiaso et al, Science 2014 |
| ATAC_seq_LSK | ATAC-seq | LSK cell | GSM1463179 | Lara-Astiaso et al, Science 2014 |
| ChIP-seq_H3K27Ac_LT_HSC | ChIP-seq | LT-HSC | GSM1441269 | Lara-Astiaso et al, Science 2014 |
| ChIP-seq_H3K27Ac_GN | ChIP-seq | Granulocytes | GSM1441277 | Lara-Astiaso et al, Science 2014 |
| ChIP-seq_H3K27Ac_Mono | ChIP-seq | Monocytes | GSM1441278 | Lara-Astiaso et al, Science 2014 |
| ChIP-seq_H3K27Ac_B | ChIP-seq | B cell | GSM1441280 | Lara-Astiaso et al, Science 2014 |
| ChIP-seq_H3K27Ac_CD4 | ChIP-seq | T cell | GSM1441281 | Lara-Astiaso et al, Science 2014 |
| ChIP-seq_H3K27Ac_CD8 | ChIP-seq | T cell | GSM1441282 | Lara-Astiaso et al, Science 2014 |

***Table S2. Sequences of Primers and sgRNAs***

| Name | Forward | Reverse | Application |
|---|---|---|---|
| HS2-1-sgRNA | CACCGAATATGTCACATTCTGTCTC | AAACGAGACAGAATGTGACATATTC | sgRNA for enhancer or promoter perturbation |
| HS2-2-sgRNA | CACCGGGACTATGGGAGGTCACTAA | AAACTTAGTGACCTCCCATAGTCCC | |
| HS2-3-sgRNA | CACCGGAAGGTTACACAGAACCAGA | AAACTCTGGTTCTGTGTAACCTTCC | |
| HS2-4-sgRNA | CACCGGCCCTGTAAGCATCCTGCTG | AAACCAGCAGGATGCTTACAGGGCC | |
| IL1RN-sgRNA1 | CACCGTGTACTCTCTGAGGTGCTC | AAACGAGCACCTCAGAGAGTACAC | |
| IL1RN-sgRNA2 | CACCGCATCAAGTCAGCCATCAGC | AAACGCTGATGGCTGACTTGATGC | |
| OCT4-sgRNA1 | CACCGACTCCACTGCACTCCAGTCT | AAACAGACTGGAGTGCAGTGGAGTC | |
| OCT4-sgRNA2 | CACCGACACCATTGCCACCACCATT | AAACAATGGTGGTGGCAATGGTGTC | |
| MYOD-sgRNA1 | CACCGGGCCCACATTCCTTTCCAG | AAACCTGGAAAGGAATGTGGGCCC | |
| MYOD-sgRNA2 | CACCGGGCTGGATTGGGTTTCCAG | AAACCTGGAAACCCAATCCAGCCC | |
| Gal4_sgRNA | CACCGAACGACTAGTTAGGCGTGTA | AAACTACACGCCTAACTAGTCGTTC | |
| Ctrl-sgRNA | CACCGTAGCGGGTAAGGATGTAGAC | AAACGTCTACATCCTTACCCGCTAC | |
| TAD1-sgRNA | CACCGTTGAACTAGTCCCGATAGGA | AAACTCCTATCGGGACTAGTTCAAC | |
| CTCF1-sgRNA | CACCGACCAGAAACACATCCCAGAT | AAACATCTGGGATGTGTTTCTGGTC | |
| CTCF2-sgRNA | CACCGTTAGACAGCAATGTTTATCC | AAACGGATAAACATTGCTGTCTAAC | |
| TAD2-sgRNA | CACCGCAGCGAGACTCCGTCGGCGT | AAACACGCCGACGGAGTCTCGCTGC | |
| HS1-sgRNA | CACCGCAATAGGTATATGAGGAGAC | AAACGTCTCCTCATATACCTATTGC | |
| HS2-sgRNA | CACCGGCCCTGTAAGCATCCTGCTG | AAACCAGCAGGATGCTTACAGGGCC | |
| HS3-sgRNA | CACCGTTTGGGAGCAGGAGTCTCTA | AAACTAGAGACTCCTGCTCCCAAAC | |
| HS4-sgRNA | CACCGCCCACTCAGCAGCTATGAGA | AAACTCTCATAGCTGCTGAGTGGGC | |
| HS5-sgRNA | CACCGTGCCCCCACCTTACAGGGAC | AAACGTCCCTGTAAGGTGGGGGCAC | |
| HS2+2.5K-sgRNA | CACCGCAAGGTGGGCAGATCACTTG | AAACCAAGTGATCTGCCCACCTTGC | |
| HS2+0.5K-sgRNA | CACCGCACAATAACTAATCATTCTA | AAACTAGAATGATTAGTTATTGTGC | |
| HS2-2.5K-sgRNA | CACCGCACTATTCCTAGCATAGTTT | AAACAAACTATGCTAGGAATAGTGC | |
| HS2-0.5K-sgRNA | CACCGAATGGTTTTGAAGCACCTGC | AAACGCAGGTGCTTCAAAACCATTC | |
| HBG-sgRNA | CACCGGCTAAACTCCACCCATGGGT | AAACACCCATGGGTGGAGTTTAGCC | |
| TAL1-Mut-sgRNA1 | CACCGagggtCACAGAAAGACGGTT | AAACAACCGTCTTTCTGTGaccctC | |
| TAL1-Mut-sgRNA2 | CACCGACAGAAAGACGGTTAGGAAA | AAACTTTCCTAACCGTCTTTCTGTC | |
| TAL1-WT-sgRNA1 | CACCGGAATGGGGTGGGGCAACCAC | AAACGTGGTTGCCCCACCCCATTCC | |
| TAL1-WT-sgRNA2 | CACCGAAAGACGTAACCCTACTTCC | AAACGGAAGTAGGGTTACGTCTTTC | |
| Spi1-P-sg1 | caccgCAAGTTCCTGATTTTATCGA | aaacTCGATAAAATCAGGAACTTGc | |
| Spi1-P-sg2 | caccgGCCCCCTCCCTTGACATTGC | aaacGCAATGTCAAGGGAGGGGGCc | |
| Spi1-P-sg3 | caccgACGGTCGTGGGTCAGACGCA | aaacTGCGTCTGACCCACGACCGTc | |
| Spi1-Enh-sg1 | caccgAGGAAGCGCCCAGTCACCAC | aaacGTGGTGACTGGGCGCTTCCTc | |
| Spi1-Enh-sg2 | caccgTGCCCAGCCTGGAGCCGCCC | aaacGGGCGGCTCCAGGCTGGGCAc | |
| Spi1-Enh-sg3 | caccgAGCTGCCCCTGTTTCCACAT | aaacATGTGGAAACAGGGGCAGCTc | |
| Cebpa-P-sg1 | caccgGTGCTAGTGGAGAGAGATCG | aaacCGATCTCTCTCCACTAGCACc | |
| Cebpa-P-sg2 | caccgACTTTCCAAGGCGGTGAGTG | aaacCACTCACCGCCTTGGAAAGTc | |
| Cebpa-Enh1-sg1 | caccgCTCCGTTCTTTTAATAGCGC | aaacGCGCTATTAAAAGAACGGAGc | |
| Cebpa-Enh1-sg2 | caccgTTGCTGCTGCTTTCTACCAC | aaacGTGGTAGAAAGCAGCAGCAAc | |
| Cebpa-Enh2-sg1 | caccgGTAATTGCCCCGGCAGAACT | aaacAGTTCTGCCGGGGCAATTACc | |
| Cebpa-Enh2-sg2 | caccgTCTGACGGGTCCTAACCTGA | aaacTCAGGTTAGGACCCGTCAGAc | |
| Cebpa-Enh3-sg1 | caccgCTTCTCATCACGGATCAAAA | aaacTTTTGATCCGTGATGAGAAGc | |
| Cebpa-Enh3-sg2 | caccgTGTTTCTGTTTTGGGGCCAC | aaacGTGGCCCCAAAACAGAAACAc | |
| Cebpa-Enh4-sg1 | caccgAAACGGGAAGTACTGAGCTA | aaacTAGCTCAGTACTTCCCGTTTc | |
| Cebpa-Enh4-sg2 | caccgGCGGGGTGAGGACATATCTC | aaacGAGATATGTCCTCACCCCGCc | |
| Gata2-P-sg1 | caccgGTCTCGGCCTCCGGCCCGCC | aaacGGCGGGCCGGAGGCCGAGACc | |
| Gata2-P-sg2 | caccgCCTGCACAGACGTGAAGCGG | aaacCCGCTTCACGTCTGTGCAGGc | |
| Gata2-Enh1-sg1 | caccgAGGTGGCCCACGGGGTGGCG | aaacCGCCACCCCGTGGGCCACCTc | |
| Gata2-Enh1-sg2 | caccgCCCTGCAGCAGGGGTTCTGC | aaacGCAGAACCCCTGCTGCAGGGc | |
| Gata2-Enh2-sg1 | caccgAACTTACGGAAACCACCTTG | aaacCAAGGTGGTTTCCGTAAGTTc | |
| Gata2-Enh2-sg2 | caccgAGGTGGATGGCCGGGCCACT | aaacAGTGGCCCGGCCATCCACCTc | |
| Gata2-Enh3-sg1 | caccgATGGCGACGCAGATAGCGCG | aaacCGCGCTATCTGCGTCGCCATc | |
| Gata2-Enh3-sg2 | caccgCCCGGAGTCTATTATACAGC | aaacGCTGTATAATAGACTCCGGGc | |
| Gata2-Enh4-sg1 | caccgACTCCGGCAGGAGATCCGAA | aaacTTCGGATCTCCTGCCGGAGTc | |
| Gata2-Enh4-sg2 | caccgGCTCTGAAAACTTGCCGGTC | aaacGACCGGCAAGTTTTCAGAGCc | |
| Gata1-P-sg1 | caccgCGGAGGGACTAGAGCCTAAA | aaacTTTAGGCTCTAGTCCCTCCGc | |
| Gata1-P-sg2 | caccgAAGGGGATCCAACAACCTGC | aaacGCAGGTTGTTGGATCCCCTTc | |
| Gata1-Enh1-sg1 | caccgATAGATAAGGGAATCAGCGG | aaacCCGCTGATTCCCTTATCTATc | |
| Gata1-Enh1-sg2 | caccgACAGAGAAGACGCTTCAACC | aaacGGTTGAAGCGTCTTCTCTGTc | |
| Gata1-Enh2-sg1 | caccgACTATTTGTGCGGAAGTCTC | aaacGAGACTTCCGCACAAATAGTc | |
| Gata1-Enh2-sg2 | caccgGTAAAGTGACAGTAAAGCGA | aaacTCGCTTTACTGTCACTTTACc | |
| Runx1-P1-sg1 | caccgGGAATGGAATTCCTCGGGTC | aaacGACCCGAGGAATTCCATTCCc | |
| Runx1-P1-sg2 | caccgGTAAAAGCACGTTCATTTCC | aaacGGAAATGAACGTGCTTTTACc | |
| Runx1-P2-sg1 | caccgGAACCACAAGTTGGGTAGCC | aaacGGCTACCCAACTTGTGGTTCc | |
| Runx1-P2-sg2 | caccgGGGTTGTGTGAAAACTTCTT | aaacAAGAAGTTTTCACACAACCCc | |
| Runx1-Enh1-sg1 | caccgGCATTATCTGCCTTCACAAG | aaacCTTGTGAAGGCAGATAATGCc | |
| Runx1-Enh1-sg2 | caccgATCCCCGCCCCAGCGACAGT | aaacACTGTCGCTGGGGCGGGGATc | |
| Runx1-Enh2-sg1 | caccgTGCAAGAGCGAGAAAACCGC | aaacGCGGTTTTCTCGCTCTTGCAc | |
| Runx1-Enh2-sg2 | caccgATGCTGACAGCCTCAGATGG | aaacCCATCTGAGGCTGTCAGCATc | |
| Runx1-Enh3-sg1 | caccgTGTGCGAGCATTGCCCCAGC | aaacGCTGGGGCAATGCTCGCACAc | |
| Runx1-Enh3-sg2 | caccgGATCTGAGTTTAGTCACTTT | aaacAAAGTGACTAAACTCAGATCc | |
| NC-sg1 | caccgGAGGTTGCAAGGGACCCGGT | aaacACCGGGTCCCTTGCAACCTCc | |
| NC-sg2 | caccgGCATTTGAGGCACACTGGAG | aaacCTCCAGTGTGCCTCAAATGCc | |
| NC-sg3 | caccgGCAGTTGCACAGATGATCAC | aaacGTGATCATCTGTGCAACTGCc | |
| NC-sg4 | caccgGGTGGTGTCGTAGTGTCAGA | aaacTCTGACACTACGACACCACCc | |
| NC-sg5 | caccgGCCTATGCTACGGACTATAG | aaacCTATAGTCCGTAGCATAGGCc | |
| NC-sg6 | caccgGGTGGCCTGGTACAGCGCAG | aaacCTGCGCTGTACCAGGCCACCc | |
| NC-sg7 | caccgGGGGCCTGACCACGATAGCA | aaacTGCTATCGTGGTCAGGCCCCc | |
| NC-sg8 | caccgGAGAGGGTTCAAAGCAGGCG | aaacCGCCTGCTTTGAACCCTCTCc | |
| NC-sg9 | caccgGCGCGAAAGGCGAGATTAAC | aaacGTTAATCTCGCCTTTCGCGCc | |
| NC-sg10 | caccgGCAGCCAAACGAAAAGACCG | aaacCGGTCTTTTCGTTTGGCTGCc | |
| IL1RN-RT | GGAATCCATGGAGGGAAGAT | TGTTCTCGCTCAGGTCAGTG | RT-qPCR primers |
| MYOD-RT | TCCCTCTTTCACGGTCTCAC | AACACCCGACTGCTGTATCC | |
| OCT4-RT | CGAAAGAGAAAGCGAACCAGTATCGAGAAC | CGTTGTGCATAGTCGCTGCTTGATCGC | |
| HBE1-RT | GCAAGAAGGTGCTGACTTCC | ACCATCACGTTACCCAGGAG | |
| HBG-RT | TGGATGATCTCAAGGGCAC | TCAGTGGTATCTGGAGGACA | |
| HBB-RT | CTGAGGAGAAGTCTGCCGTTA | AGCATCAGGAGTGGACAGAT | |
| GAPDH-RT | ACCCAGAAGACTGTGGATGG | TTCAGCTCAGGGATGACCTT | |
| TAL1-RT | GTGCAGCTGAGTCCTCCCGC | CCGGCTCCCCAAAGAACCCG | |
| Spi1-RT | TACCAACGTCCAATGCATGACTAC | GAGTATCGAGGACGTGCATCTGTT | |
| Runx1-RT | CCAGCAAGCTGAGGAGCGGCG | TGACGGTGACCAGAGTG | |
| Gata1-RT | CAGAACCGGCCTCTCATCC | TAGTGCATTGGGTGCCTGC | |
| Gata2-RT | GCAGAGAAGCAAGGCTCGC | CAGTTGACACACTCCCGGC | |
| Cebpa-1-RT | GCGGGAACGCAACAACATC | GTCACTGGTCAACTCCAGCAC | |
| gDNA_sgRNA_NGS_LCreadoutPRC1-F | AATGGACTATCATATGCTTACCGTAACTTGAAAGTATTTCG | AAAAAAAgcaccgactcggtgccacttggccCTGCAGACA | sgRNA pooled screen primers with barcodes and staggers |
| sgRNApool_NGS_F1 | AATGATACGGCGACCACCGAGATCTACACTCTTTCCCTACACGACGCTCTTCCGATCTtAAGTAGAGtcttgtggaaaggacgaaacaccg | | |
| sgRNApool_NGS_F2 | AATGATACGGCGACCACCGAGATCTACACTCTTTCCCTACACGACGCTCTTCCGATCTatACACGATCtcttgtggaaaggacgaaacaccg | | |
| sgRNApool_NGS_F3 | AATGATACGGCGACCACCGAGATCTACACTCTTTCCCTACACGACGCTCTTCCGATCTgatCGCGCGGTtcttgtggaaaggacgaaacaccg | | |
| sgRNApool_NGS_F4 | AATGATACGGCGACCACCGAGATCTACACTCTTTCCCTACACGACGCTCTTCCGATCTcgatCATGATCGtcttgtggaaaggacgaaacaccg | | |
| sgRNApool_NGS_F5 | AATGATACGGCGACCACCGAGATCTACACTCTTTCCCTACACGACGCTCTTCCGATCTtcgatCGTTACCAtcttgtggaaaggacgaaacaccg | | |
| sgRNApool_NGS_F6 | AATGATACGGCGACCACCGAGATCTACACTCTTTCCCTACACGACGCTCTTCCGATCTatcgatTCCTTGGTtcttgtggaaaggacgaaacaccg | | |
| sgRNApool_NGS_F7 | AATGATACGGCGACCACCGAGATCTACACTCTTTCCCTACACGACGCTCTTCCGATCTgatcgatAACGCATTtcttgtggaaaggacgaaacaccg | | |
| sgRNApool_NGS_F8 | AATGATACGGCGACCACCGAGATCTACACTCTTTCCCTACACGACGCTCTTCCGATCTcgatcgatACAGGTATtcttgtggaaaggacgaaacaccg | | |
| sgRNApool_NGS_F9 | AATGATACGGCGACCACCGAGATCTACACTCTTTCCCTACACGACGCTCTTCCGATCTacgatcgatAGGTAAGGtcttgtggaaaggacgaaacaccg | | |
| sgRNApool_NGS_F10 | AATGATACGGCGACCACCGAGATCTACACTCTTTCCCTACACGACGCTCTTCCGATCTtAACAATGGtcttgtggaaaggacgaaacaccg | | |
| sgRNApool_NGS_F11 | AATGATACGGCGACCACCGAGATCTACACTCTTTCCCTACACGACGCTCTTCCGATCTatACTGTATCtcttgtggaaaggacgaaacaccg | | |
| sgRNApool_NGS_F12 | AATGATACGGCGACCACCGAGATCTACACTCTTTCCCTACACGACGCTCTTCCGATCTgatAGGTCGCAtcttgtggaaaggacgaaacaccg | | |
| PMLS_POOL_NGS_R1 | CAAGCAGAAGACGGCATACGAGATAAGTAGAGGTGACTGGAGTTCAGACGTGTGCTCTTCCGATCTtTCGGTGCCACTTGGCCCTGCAGACA | | |
| PMLS_POOL_NGS_R2 | CAAGCAGAAGACGGCATACGAGATACACGATCGTGACTGGAGTTCAGACGTGTGCTCTTCCGATCTatTCGGTGCCACTTGGCCCTGCAGACA | | |
| PMLS_POOL_NGS_R3 | CAAGCAGAAGACGGCATACGAGATCGCGCGGTGTGACTGGAGTTCAGACGTGTGCTCTTCCGATCTgatTCGGTGCCACTTGGCCCTGCAGACA | | |
| PMLS_POOL_NGS_R4 | CAAGCAGAAGACGGCATACGAGATCATGATCGGTGACTGGAGTTCAGACGTGTGCTCTTCCGATCTcgatTCGGTGCCACTTGGCCCTGCAGACA | | |
| PMLS_POOL_NGS_R5 | CAAGCAGAAGACGGCATACGAGATCGTTACCAGTGACTGGAGTTCAGACGTGTGCTCTTCCGATCTcgatcTCGGTGCCACTTGGCCCTGCAGACA | | |
| PMLS_POOL_NGS_R6 | CAAGCAGAAGACGGCATACGAGATTCCTTGGTGTGACTGGAGTTCAGACGTGTGCTCTTCCGATCTtTCGGTGCCACTTGGCCCTGCAGACA | | |
| rtTA-A | AAAGTCGCTCTGAGTTGTTAT | | mouse genotyping primers |
| rtTA-B | GCGAAGAGTTTGTCCTCAACC | | |
| rtTA-C | GGAGCGGGAGAAATGGATATG | | |
| dCas9-KRAB-A | GCACAGCATTGCGGACATGC | | |
| dCas9-KRAB-B | CCCTCCATGTGTGACCAAGG | | |
| dCas9-KRAB-C | GCAGAAGCGCGGCCGTCTGG | | |
